# Dimensionality reduction methods for extracting functional networks from large-scale CRISPR screens

**DOI:** 10.1101/2023.02.22.529573

**Authors:** Arshia Zernab Hassan, Henry N. Ward, Mahfuzur Rahman, Maximilian Billmann, Yoonkyu Lee, Chad L. Myers

**Affiliations:** Department of Computer Science and Engineering, University of Minnesota – Twin Cities, Minneapolis, Minnesota, USA; Bioinformatics and Computational Biology Graduate Program, University of Minnesota – Twin Cities, Minneapolis, Minnesota, USA; Institute of Human Genetics, University of Bonn, School of Medicine and University Hospital Bonn, Bonn, Germany

**Keywords:** Gene co-essentiality network, Normalization, Unsupervised dimensionality reduction, Auto-encoder, Robust principal component analysis

## Abstract

CRISPR-Cas9 screens facilitate the discovery of gene functional relationships and phenotype-specific dependencies. The Cancer Dependency Map (DepMap) is the largest compendium of whole-genome CRISPR screens aimed at identifying cancer-specific genetic dependencies across human cell lines. A mitochondria-associated bias has been previously reported to mask signals for genes involved in other functions, and thus, methods for normalizing this dominant signal to improve co-essentiality networks are of interest. In this study, we explore three unsupervised dimensionality reduction methods - autoencoders, robust, and classical principal component analyses (PCA) - for normalizing the DepMap to improve functional networks extracted from these data. We propose a novel “onion” normalization technique to combine several normalized data layers into a single network. Benchmarking analyses reveal that robust PCA combined with onion normalization outperforms existing methods for normalizing the DepMap. Our work demonstrates the value of removing low-dimensional signals from the DepMap before constructing functional gene networks and provides generalizable dimensionality reduction-based normalization tools.

## Introduction

Deciphering the functional relationships among genes is imperative for understanding the mechanism of diseases with genetic components. Whole-genome CRISPR screening is one state-of-the-art method for identifying phenotype-specific genetic dependencies for diseases like cancer (Shalem, et al., 2014; Wang, Wei, Sabatini, & Lander, 2014; Tsherniak, et al., 2017). In addition to identifying cancer-specific dependencies, high-throughput data generated from whole-genome CRISPR screens can be mined to map functional relationships between genes (Pan, et al., 2018; Boyle, Pritchard, & Greenleaf, 2018; Wainberg, et al., 2021; Kim, et al., 2019; Buphamalai, Kokotovic, Nagy, & Menche, 2021). Therefore, the development of novel algorithms to process, normalize and mine whole-genome CRISPR screening data could prove particularly fruitful for identifying such functional relationships.

Most CRISPR screens use CRISPR-Cas9 guides to introduce targeted knockouts across the vast majority of the human genome in human cell culture. In brief, the workflow for a typical screen involves the infection of human cell culture with a lentiviral vector containing a library of ∼70,000 guide (g)RNAs targeting around 18,000 genes. After passaging the cell population over several days, sequencing performed at various timepoints measures the dropout of gRNAs from the population. At the end of the experiment, computational analyses are performed to quantify observed fitness effects relative to controls, such as known non-essential guides or screens performed in wildtype cells. Current experimental techniques for performing whole-genome CRISPR screens are perhaps best exemplified by the Cancer Dependency Map (DepMap) project’s efforts to discover genetic dependencies across human cell lines (Tsherniak, et al., 2017; Meyers, et al., 2017; Behan, et al., 2019; Dempster, et al., 2019; Pacini, et al., 2021; Dharia, et al., 2021). As of the 22Q4 version, the Cancer Dependency Map project has performed such CRISPR screens to identify cancer-specific genetic dependencies across 1,078 cell lines (Data ref: (Broad, 2022)).

In addition to directly identifying cancer-specific genetic dependencies, co-essentiality between genes can be measured and used to group genes into functional modules by measuring correlations between CERES scores in the DepMap - a type of analysis pioneered in the yeast genetic interaction research community (Baryshnikova, et al., 2010; Costanzo, et al., 2016). Indeed, this profile similarity analysis has been directly applied to the DepMap dataset to reveal functional similarities between human genes (Pan, et al., 2018; Boyle, Pritchard, & Greenleaf, 2018; Wainberg, et al., 2021; Kim, et al., 2019; Buphamalai, Kokotovic, Nagy, & Menche, 2021; Gheorghe & Hart, 2022). However, previous research has posited that profile similarities in the DepMap are confounded by technical variation unrelated to the cancer-specific phenotypes of interest (Rahman, et al., 2021).

To address this problem, two methods for computationally enhancing cancer-specific signals and identifying the source of variation attributable to technical factors from the DepMap have been proposed. Boyle et al. proposed to remove principal components derived from olfactory receptor gene profiles, which are assumed to contain variation irrelevant to cancer-specific dependencies, from the data (Boyle, Pritchard, & Greenleaf, 2018). A separate method proposed by Wainberg et al. to enhance signals within the DepMap applied generalized least squares (GLS) to account for dependence among cell lines (Wainberg, et al., 2021). Our own functional evaluation of DepMap profiles using external gold-standards such as CORUM (Comprehensive Resource of Mammalian protein complex) protein co-complex annotations revealed substantial bias related to mitochondrial complexes, which dominate typical correlation analyses of DepMap profiles (Rahman, et al., 2021). These signals are highly biologically relevant, but their dominance may eclipse contributions of genes in smaller complexes, which also represent cancer-specific dependencies. Because these existing normalization techniques have shown mixed results for boosting signal within smaller and non-mitochondrial complexes, in this study, we explore the use of unsupervised dimensionality reduction approaches for normalizing the DepMap dataset.

We explore classical principal component analysis (PCA) as well as two state-of-the-art dimensionality reduction normalization methods’ abilities to boost the signal of cancer-specific dependencies and remove mitochondrial signal from the DepMap (Wold, Esbensen, & Geladi, 1987). We also propose a novel method for integrating signal across different levels of normalized data. Specifically, we apply a variant of PCA called robust PCA (RPCA) as well as autoencoder neural networks (AE) to learn and remove confounding low-dimensional signal from the DepMap (Candès, Li, Ma, & Wright, 2011; Hinton & Salakhutdinov, 2006). In addition, we propose a novel method named “onion” normalization as a general-purpose technique for integrating multiple layers of normalized data across different hyperparameter values into a single normalized dataset. We apply onion normalization using either PCA-normalized, RPCA-normalized or AE-normalized data as input. Our benchmarking analyses of the normalized versions of the DepMap demonstrate that, while autoencoder normalization most efficiently captures and removes mitochondrial-associated signal from the DepMap, aggregating signals across different layers with onion normalization applied to RPCA-normalized networks is most effective at enhancing functional relationships between genes in the DepMap dataset.

## Results

### Removing low-dimensional signal from the DepMap boosts the performance of non-mitochondrial complexes

Dimensionality reduction techniques aim to transform a high-dimensional dataset into a low-dimensional one, and although they are typically applied under the assumption that low-dimensional signal is desirable (Way & Greene, 2018; Lotfollahi, Wolf, & Theis, 2019; Ding, Condon, & Shah, 2018; Lopez, Regier, Cole, Jordan, & Yosef, 2018; Sun, Zhu, Ma, & Zhou, 2019), we flip that assumption in order to normalize DepMap data. We posit that two properties of the DepMap hold: we assume that true genetic dependencies are rare, based on estimations from large-scale yeast genetic interaction studies (Costanzo, et al., 2016), and we assume that dominant low-dimensional signal in the DepMap is likely to represent mitochondrial-associated bias that is plausibly driven by technical variation or non-specific biological variation. For example, in the only genome-wide study of GIs to date, it was estimated that an average gene interacts with others roughly 3% of the time (Costanzo, et al., 2016). Therefore, instead of assuming that low-dimensional representations of DepMap data are desirable for data mining and visualization purposes, we instead propose to capture and remove that dominant signal from the DepMap (Figure 1A, 1B). We applied multiple dimensionality reduction methods to the DepMap to accomplish this goal, beginning with classical PCA normalization. To explore the extent to which normalization improves the detection of functional relationships between genes and removes mitochondrial bias from the DepMap, we applied benchmarking analyses with a software package developed for this purpose called FLEX (Rahman, et al., 2021).

**Figure 1:**
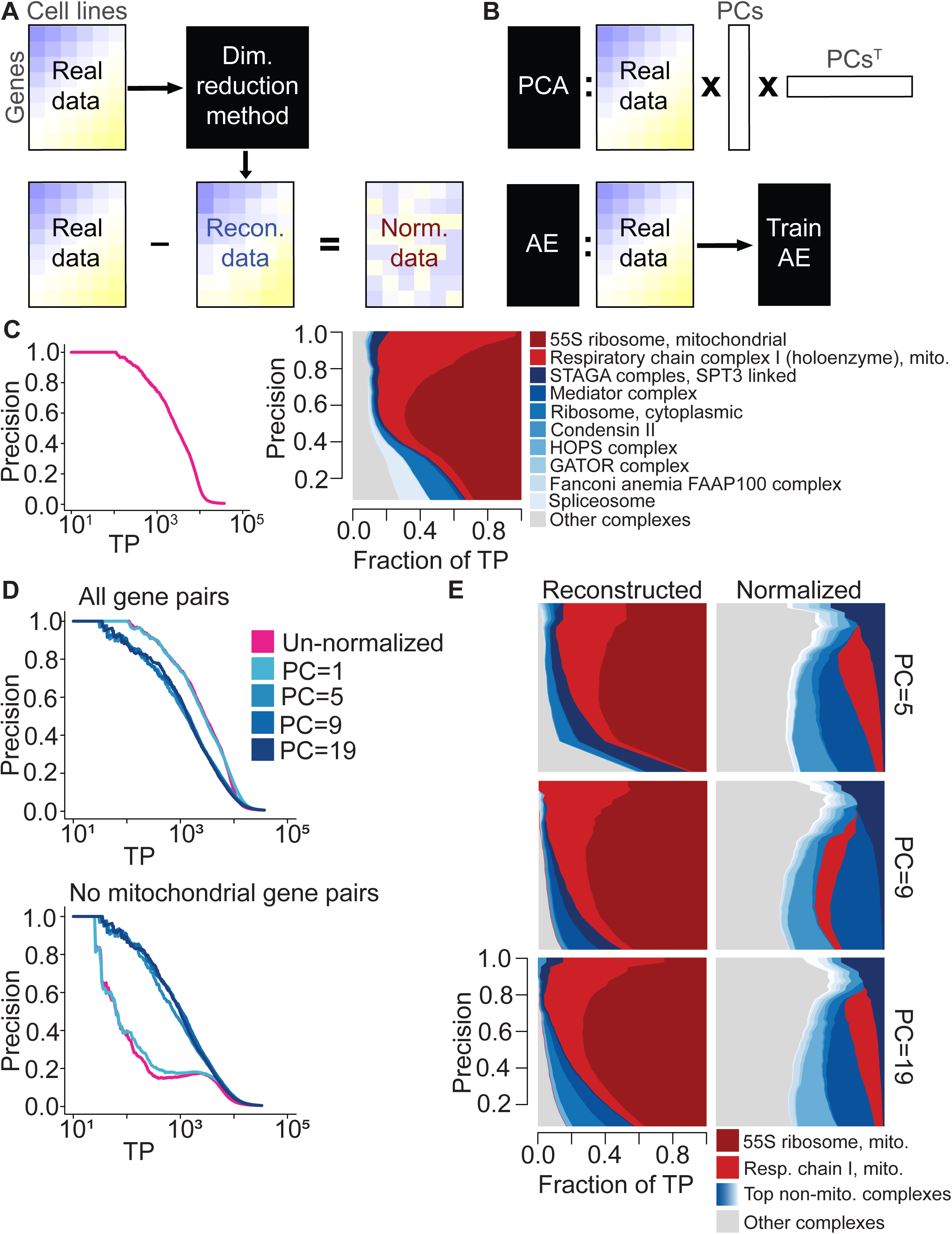
Normalization schematic and exploration of mitochondrial bias within the DepMap for different numbers of PCs removed. **A**, A dimensionality reduction method is applied to the original DepMap data to extract a low-dimensional representation of the data. Reconstructed data is generated from that, which is subtracted from the original DepMap to normalize it. **B**, (Top) PCA generates reconstructed DepMap data by multiplying the DepMap against selected PCs derived from it and the transpose of those PCs. (Bottom) Autoencoders generate reconstructed data post-training by passing in the original DepMap as input. **C**, (Left) Precision-recall (PR) performance of un-normalized DepMap data evaluated against CORUM protein complexes. The x-axis depicts the log-scaled number of true-positives (TPs). (Right) Contribution diversity plot of CORUM complexes in un-normalized DepMap data. This plot is constructed by sliding a precision cutoff from high to low (indicated by the y-axis), and at each point, plotting a stacked bar plot across the x-axis at that point reflecting the breakdown of complex membership of the TP pairs identified at that threshold. The top ten contributing complexes are listed in the legend, with the light gray category representing all complexes represented at lower frequency. **D**, (Top) Precision-recall (PR) performance of PCA-normalized data with the first 5, 9, and 19 principal components removed evaluated against CORUM protein complexes. (Bottom) PR performance with mitochondrial gene pairs removed from evaluation. **E**, The contribution diversity plots depict CORUM complex contributions in PCA-reconstructed data and PCA-normalized data for the first 5, 9 and 19 principal components.

Benchmarking analyses with FLEX based on the CORUM protein complex standard reveal the extent of mitochondrial dominance in the DepMap for both the original dataset and all normalized versions (Giurgiu, et al., 2019). To summarize this benchmarking process, a gene-level similarity matrix is created from the per-gene dependency scores by calculating Pearson correlation coefficients (PCCs) between all pairs of genes. Taking these similarity scores and a set of gold standard co-annotations for genes as input, FLEX generates precision-recall curves (PR curves) that measure how many true positive gene pairs in the gold standard set are recapitulated by PCCs taken at different similarity thresholds. More detailed information such as which complexes drive the performance of PR curves are also output by FLEX and are illustrated graphically by diversity plots. To interpret these plots, a visually larger area corresponds to more contribution to the overall PR curve from a complex at the corresponding precision threshold. An examination of the original DepMap’s CORUM PR curve performance alongside a diversity plot reveals that most performance in the PR curve is driven by two mitochondria-related complexes - the 55S ribosome and respiratory chain complexes (Figure 1C). Therefore, to ascertain how much signal the DepMap contains for all other protein complexes, we generated PR curves that exclude a set of mitochondrial genes and observed a drastic but expected drop in overall performance (Figure 1D, Methods).

As a reference dimensionality reduction technique, we first examined the extent to which classical PCA captures mitochondrial signal and boosts signal from other complexes post-normalization. In the PCA-normalization approach, PCA is first applied to gene perturbation profiles to capture low-dimensional signal. Then, the original dataset is projected onto a subset of the strongest PCs to generate a “reconstructed” version of the DepMap. Directly subtracting the reconstructed DepMap from the original DepMap produces a PCA-normalized version of the DepMap that does not contain the signal from the selected PCs.

While PCA-normalization has already been applied to DepMap versions starting from 2019 Q3 to remove several principal components, this is insufficient to reduce the mitochondrial dominance of the dataset or to boost signal within smaller complexes (Data ref: (Broad, 2019)). Repeating analyses detailed in Rahman et al., which analyzed the 18Q3 and 19Q2 versions of the DepMap, for the 20Q2 version, which is used for all analyses in this manuscript, reveals that co-dependency profiles are still dominated by mitochondrial signals (Rahman, et al., 2021) (Data ref: (Broad, 2018; Broad, 2019; Broad, 2020)). In addition to removing this signal, successful normalization methods have the potential to uncover relationships masked by this signal, which can be measured by observing boosts in the performance of smaller complexes in terms of their contributions to CORUM PR curves.

Surprisingly, removing a large number of principal components from the DepMap improves the dataset’s ability to capture signal within non-mitochondrial complexes (Figure 1D, 1E). We applied PCA-normalization to the DepMap 20Q2 dataset and removed a varying number of principal components - either 1, 5, 9 or 19 (Broad, 2020). In addition to generating standard CORUM PR curves with FLEX as described above, to measure the ability of each dataset to recover signal within non-mitochondrial complexes, we also generated PR curves where mitochondrial gene pairs were removed as positive examples from the CORUM standard (Figure 1D). While this only affects gene pairs where both genes are members of a set of 1,266 genes (see Methods), these mitochondrial-attenuated PR curves nevertheless reveal that removing 5 or more principal components boosts signal for non-mitochondrial complexes compared to the original DepMap. Diversity plots generated with FLEX confirm this observation (Figure 1E, Figure S9, Figure S10). We conclude that functional signal for most protein complexes remains and even improves while mitochondrial signal in the DepMap decreases after removing many principal components. These observations suggest that the strongest low-dimensional components of the DepMap are likely to represent technical variation, or at least non-specific variation that clouds more specific functional information, and that removing a large number of low-dimensional components is valuable in measuring functional relationships.

In the following section, we introduce two state-of-the-art dimensionality reduction techniques for normalizing the DepMap before characterizing their ability to both reduce the dominance of mitochondrial-associated signal and boost the performance of smaller complexes.

### Autoencoder and robust PCA normalization robustly capture and remove technical variation from the DepMap

Autoencoders are a type of deep neural network method designed for unsupervised dimensionality reduction (Hinton & Salakhutdinov, 2006). They function by optimizing the generation of reconstructed profiles that are similar to a training dataset after passing the training data through a neural network constructed in an “hourglass” shape. A crucial parameter of autoencoders is the latent space size, referred to as *LS* throughout, which is the number of nodes contained in the bottleneck layer at the center of the hourglass.

Strikingly, our analysis shows that deep convolutional autoencoders trained with a single-dimensional latent space can both generate realistic reconstructed profiles as well as capture and remove the majority of signal contributed by mitochondrial complexes in the DepMap. Similar to PCA-normalization, after training the autoencoder and observing high gene-wise correlations between reconstructed profiles and the original profiles, we created AE-normalized data by directly subtracting the reconstructed matrix from the original data, thereby removing the low-dimensional signal. FLEX benchmarking shows that AE-normalized data for *LS* = 1, where the bottleneck layer consists of only a single node, strongly reduces the dominance of mitochondrial complexes while boosting the signal of non-mitochondrial complexes (Figure 2A, Figure S13, Figure S14), similar to PCA-normalization with many principal components. This provides evidence that the mitochondrial signal in the DepMap is low-dimensional and can be captured efficiently with an autoencoder model.

**Figure 2:**
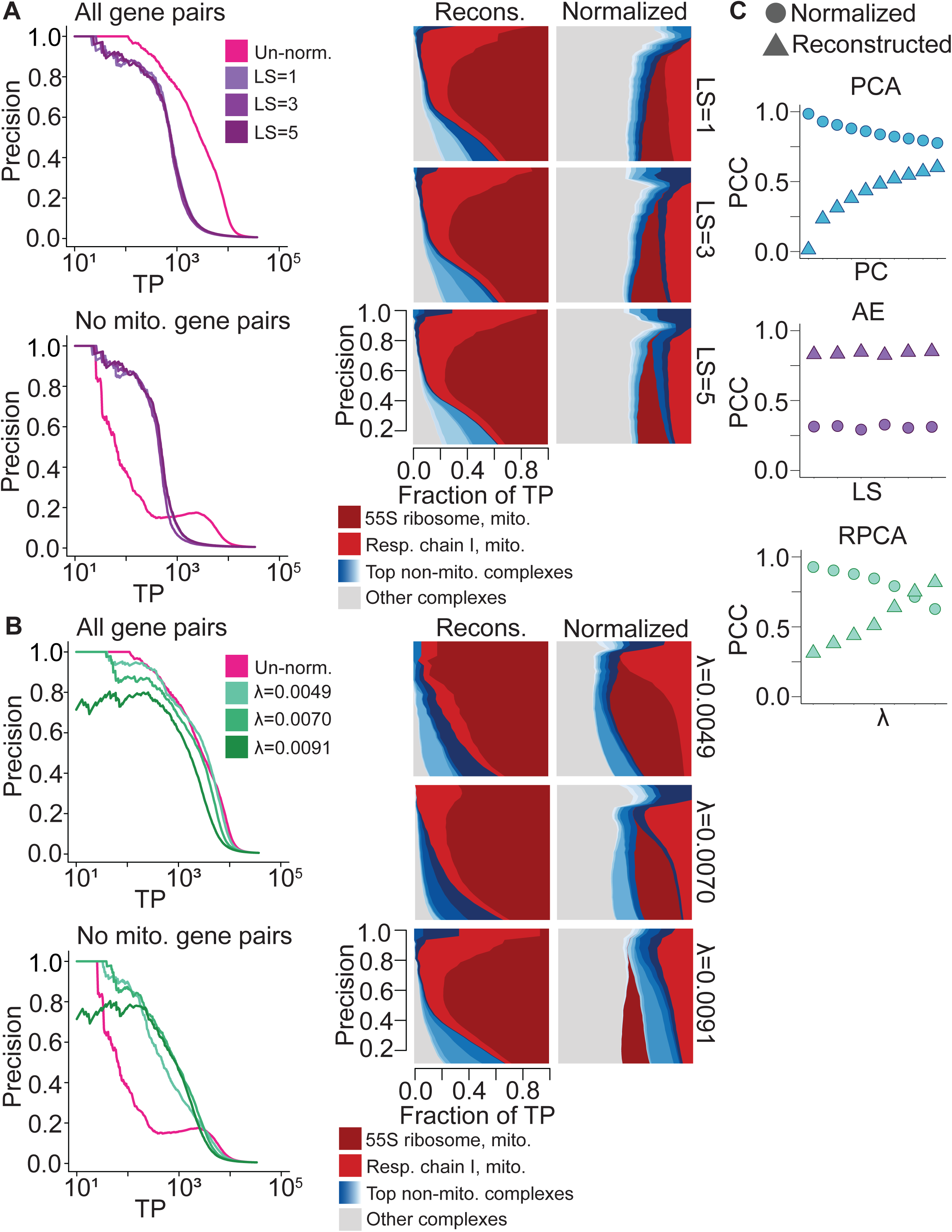
Exploration of PCA, RPCA and AE normalization across hyperparameters. **A**, (Left) Precision-recall (PR) performance of AE-normalized data generated with latent space sizes 1, 3 and 5 evaluated against CORUM protein complexes. (Right) Corresponding contribution diversity plots depicting CORUM complex contributions from AE-reconstructed and AE-normalized data. **B**, (Left) PR performance of RPCA-normalized data generated with *λ* set to 0.0049, 0.007 and 0.0091 evaluated against CORUM protein complexes. (Right) Corresponding contribution diversity plots illustrating complex contributions in RPCA-reconstructed and RPCA-normalized data. **C**, Scatter plot of Pearson correlation coefficients between un-normalized data and reconstructed data as well as between un-normalized data and normalized data generated by PCA, AE and RPCA normalization. Y-axis contains Pearson correlation coefficient values, and the X-axis contains the number of removed principal components (first 1, 3, 5, 7, 9, 11, 13, 15, 17, 19) for PCA-normalization, latent space sizes (1, 2, 3, 4, 5, 10) for AE-normalization and *λ* (approximately 0.0049, 0.0056, 0.0063, 0.007, 0.0077, 0.0084, 0.0091) for RPCA-normalization.

The second normalization technique that we apply to the DepMap is robust principal component analysis (Candès, Li, Ma, & Wright, 2011). RPCA, a modified version of PCA, is an unsupervised technique used to decompose a matrix into two components: a low-dimensional component and a sparse component, which are assumed to be superimposed. In this context, we expect the low-rank component to capture technical or non-specific biological variation and the sparse component to capture true genetic dependencies. Indeed, when we applied RPCA to the DepMap, it separated most of the dominant mitochondrial signals into the low-rank component (the “reconstructed” dataset) while the sparse component retained high-quality information about other functional relationships (the “normalized” dataset; Figure 2B, Figure S11, Figure S12). Dialing *λ*, a hyperparameter of RPCA, controls the dimensionality of the low-rank component, with smaller values increasing the dimensionality of the low-rank component.

Autoencoder and RPCA normalization consistently generated realistic reconstructed data and boosted the performance of smaller complexes across different values of *LS* and *λ*, respectively. Autoencoders trained with different values of *LS* generated reconstructed data with similarly high Pearson correlations to the original DepMap dataset, consistent with the observation that an autoencoder with a bottleneck layer consisting of a single layer efficiently captures most mitochondrial signal in the DepMap. On the other hand, RPCA runs for larger values of *λ* resulted in reconstructed datasets with drastically improved correlation to the original DepMap, similar to the behavior of classical PCA (Figure 2C). Both autoencoder and RPCA normalization contributed consistent performance increases for non-mitochondrial complexes within CORUM PR curves (Figure 2A, 2B).

Interestingly, closer examination of the complexes with improved signal revealed that different complexes peaked in terms of performance at different hyperparameter settings for all methods (Figure S1, Figure S2, Figure S3). Therefore, we sought to apply a method that could integrate normalized datasets across several different hyperparameter choices to maximize performance in detecting varied functional relationships in normalized data.

### Onion normalization integrates normalized data across hyperparameter values

The final normalization technique we propose directly addresses this problem and involves the integration of several “layers” of normalized data - where different layers are versions of the DepMap normalized based on specific hyperparameter values, such as AE-normalized data for varying values of *LS* - in order to assimilate rare signals that may not be present in all layers of the data. The core assumption of “onion” normalization, which is supported by our previous analyses of both PCA-normalized and AE-normalized data, is that dialing the parameter values of a specific normalization method yields normalized gene effect scores containing information specific to individual layers as well as information common to multiple layers. As a result, similarity networks created using differently-normalized networks may convey information with substantial variation, with each one capturing informative relationships between genes. Thus, to summarize the diverse information contained in separate layers of normalized data and to avoid computational and analytical redundancy, “onion” normalization aims to incorporate many different layers of normalized data into a single network.

We used a previously-published, unsupervised technique called similarity network fusion (SNF) to perform this integration (Wang, et al., 2014). SNF operates by integrating several similarity networks using a network fusion technique based on multiview learning that considers the neighborhood and sparsity information of individual networks, which can integrate networks with subtle differences in an unbiased manner.

A key strength of onion normalization is that any effective dimensionality reduction method can be employed in the normalization step to generate different layers of the “onion.” The similarity network layers to be fused are created from the same data normalized by varying key parameters of the chosen normalization method. For this study, we compared onion normalization applying PCA-normalization with varying numbers of PCs (PCO), autoencoder normalization with varying latent space sizes (AEO), and RPCA normalization with varying lambda values (RPCO) (Figure 3A).

**Figure 3:**
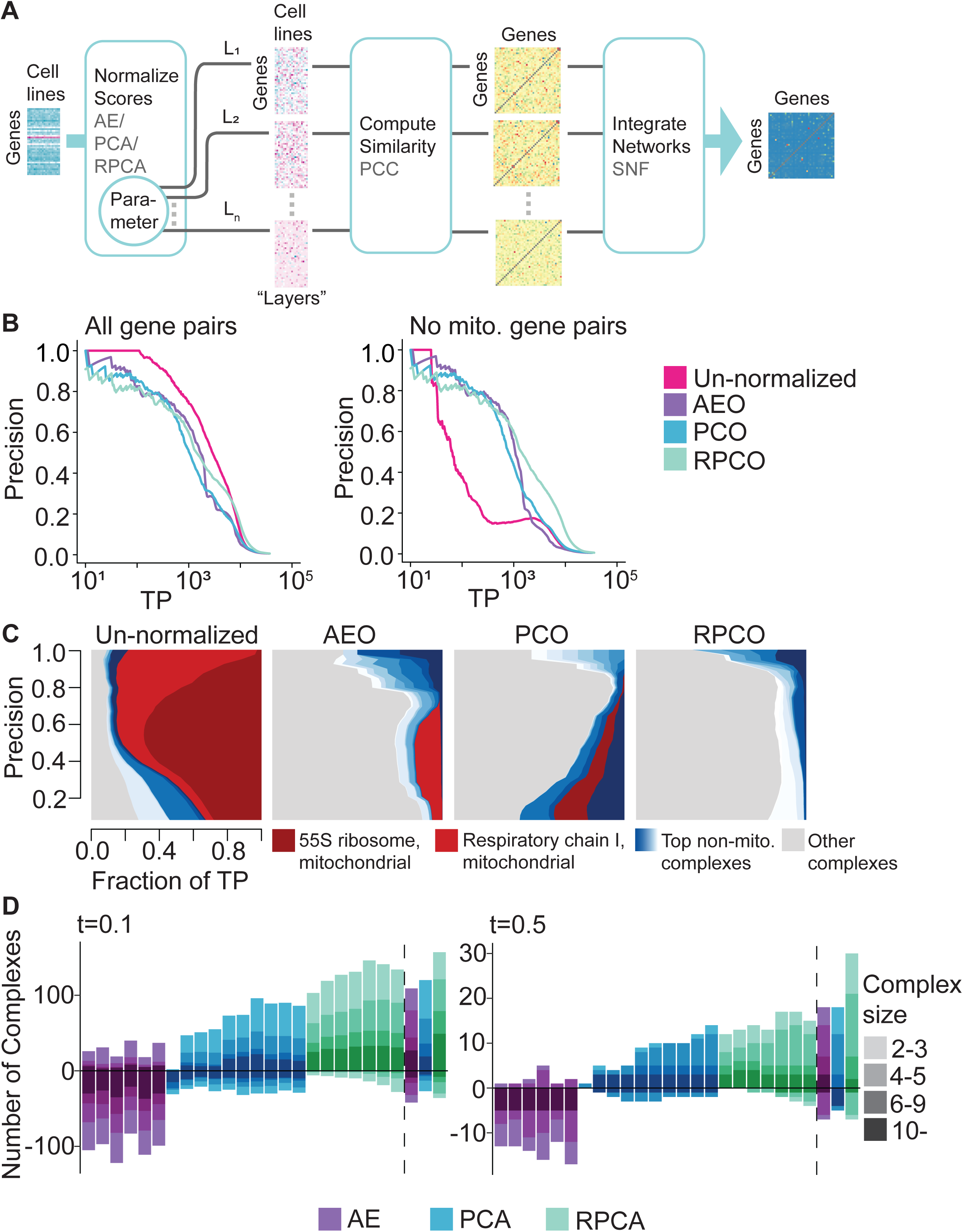
Onion normalization and benchmarking for different normalization techniques as input. **A**, Schematic of onion normalization. Pearson correlation coefficients are computed from data normalized with a chosen technique - autoencoders (AE), PCA or robust PCA (RPCA) - for different choices of hyperparameters, which are then integrated with similarity network fusion (SNF). **B**, FLEX precision-recall (PR) performance of original DepMap CERES scores against onion normalization with AE (AEO), PCA (PCO) or RPCA (RPCO) as input for CORUM protein complexes as the standard. (Left) All CORUM co-complex pairs as true positives. (Right) Mitochondrial gene pairs are removed from true positives. **C**, Contribution diversity of CORUM complexes for the original DepMap, AEO, PCO and RPCO data. Fractions of predicted true positives (TP) from different complexes are plotted at various precision levels on the y-axis. **D**, Number of complexes for which area under the PR curve (AUPRC) values increase and decrease with respect to chosen AUPRC thresholds due to normalization as compared to un-normalized data. The bars on the left side of the dotted line correspond to AE-normalized layers (latent space size = 1, 2, 3, 4, 5, 10), PCA-normalized layers (first 1, 3, 5, 7, 9, 1, 13, 15, 17, 19 principal components removed) and RPCA-layers (*λ* ~= 0.0049, 0.0056, 0.0063, 0.007, 0.0077, 0.0084, 0.0091). The bars on the right side of the dotted line correspond to SNF integrated data of the respective layers for all three methods. The color gradient for each method represents four bins with complexes containing 2 to 3 genes, 4 to 5 genes, 6 to 9 genes, and 10 or more genes. (Left) *t* = 0.1. (Right) *t* = 0.5.

FLEX benchmarking reveals that onion normalization improves performance compared to individual layers of normalized data for all normalization methods, with RPCO normalization showing the strongest performance of the three approaches (Figure 3B, 3D). Mitochondrial-attenuated PR curves reveal a substantial performance benefit for all onion-normalized datasets compared to the original DepMap. Moreover, due to improved performance for boosting weaker signal later in the PR curve (i.e., at thresholds corresponding to higher recall), RPCO outperforms both PCO and AEO (Figure 3B). Diversity plots of CORUM PR curves suggest that RPCO-normalization greatly reduces the mitochondrial dominance observed in the original DepMap dataset (Figures 3C, Figure S15). However, a closer analysis of the complexes driving the RPCO diversity plot reveals that, in addition to a partial reduction of mitochondrial-associated signal, signal within non-mitochondrial complexes is boosted such that the ten complexes driving PR curve performance no longer include mitochondrial-associated complexes. Thus, rather than normalizing mitochondrial signal entirely out of the DepMap, RPCO normalization instead boosts signal within smaller, non-mitochondrial complexes such that the strongest gene-gene similarities are no longer dominated by mitochondria-related genes. All onion-normalized datasets also outperform their individual normalized layers for boosting signal within smaller complexes (Figure 3D).

A detailed analysis of complexes with boosted signal across normalization techniques shows that RPCO normalization best improves the signal contained in complexes with low signal in the original DepMap. We plotted the number of complexes with strongly boosted or weakened signal, defined as those with AUPRCs that differ by “small” magnitudes of 0.1 or “large” magnitudes of 0.5 in normalized data compared to the original DepMap, and binned those across complex size for all normalization techniques (Figure 3D). This analysis shows that integration with onion normalization, especially with RPCA, outperforms all individually-normalized layers at boosting the signal contained across complexes of different sizes. For example, even though autoencoder normalization efficiently removes mitochondrial signal, it also removes signal from many non-mitochondrial complexes - a drawback rescued by integration with onion normalization.

Similar benchmarking analyses show that RPCO and AEO-normalization outperform the GLS normalization technique proposed by Wainberg et al. and the olfactory receptor normalization (OLF) technique proposed by Boyle et al. (Wainberg, et al., 2021; Boyle, Pritchard, & Greenleaf, 2018). Mitochondrial-attenuated PR curves show improved performance of RPCO over AEO and GLS, which perform similarly (Figure 4A), while diversity plots reveal that both AEO and RPCO reduce mitochondrial-associated signal more distinctly than GLS (Figure 4B, Figure S16). Plotting per-complex AUPRC values based on the magnitude of differences compared to un-normalized data for all methods details a similar pattern for magnitude thresholds of 0.1 and 0.5, where RPCO performs best and AEO and GLS perform similarly (Figure 4C). For the complexes with the most pronounced difference between unnormalized and normalized data at a magnitude threshold of 0.7 AUPRC, both RPCO and AEO perform similarly and substantially outperform GLS. Across all evaluations, OLF normalization does not substantially reduce mitochondrial signal or boost signal contained within non-mitochondrial complexes compared to the other three methods.

**Figure 4:**
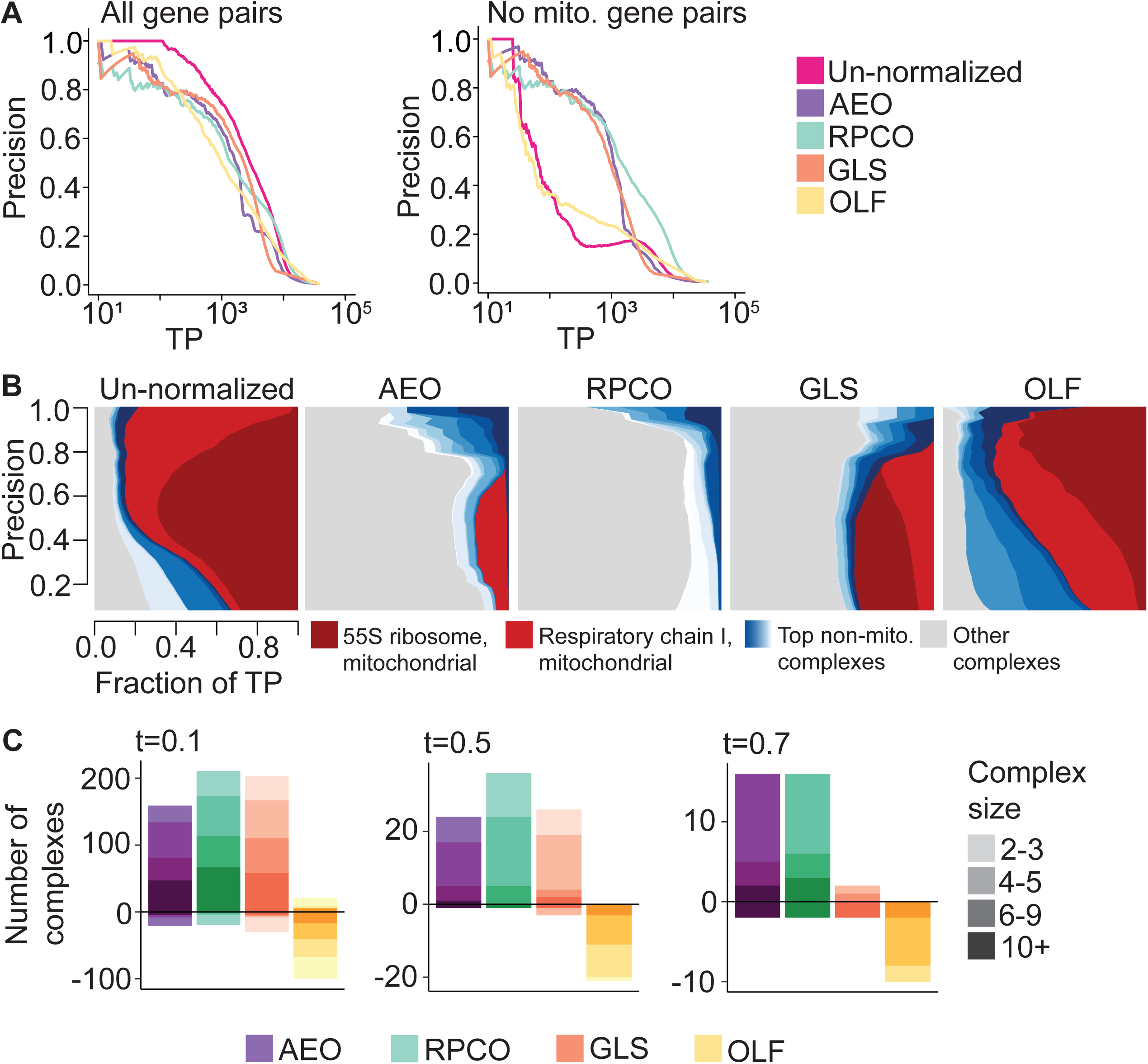
Onion normalization and benchmarking for different normalization techniques as input. **A**, FLEX precision-recall (PR) performance of original DepMap CERES scores against onion normalization with AE (AEO), onion normalization with robust PCA (RPCO), generalized least squares (GLS) normalization from in Wainberg et al. (Wainberg, et al., 2021), and olfactory receptor (OLF) normalization from Boyle et al. (Boyle, Pritchard, & Greenleaf, 2018) as input for CORUM protein complexes as the standard. (Left) All CORUM co-complex pairs as true positives. (Right) Mitochondrial gene pairs are removed from true positives. **B**, Contribution diversity of CORUM complexes for the original DepMap, AEO, RPCO, GLS and OLF data. Fractions of predicted true-positives (TP) from different complexes are plotted at various precision levels on the y-axis. **C**, Number of complexes for which area under the PR curve (AUPRC) values increase and decrease with respect to chosen AUPRC thresholds due to normalization as compared to un-normalized data for AEO, RPCO, GLS and OLF data. The color gradient for each method represents four bins with complexes containing 2 to 3 genes, 4 to 5 genes, 6 to 9 genes, and 10 or more genes. (Left) *t* = 0.1. (Middle) *t* = 0.5. (Right) *t* = 0.7.

### Network analysis of onion-normalized DepMap data uncovers biologically relevant clusters

To visually examine functional relationships between genes pre- and post-RPCO normalization and the expected reduction in mitochondrial signal, we created correlation networks for both versions of the DepMap in Cytoscape version 3.7.2 (Shannon, et al., 2003) using the yFiles organic layout algorithm. We performed this for five, ten, and fifteen thousand of the top-ranked edges sorted in decreasing order of correlations for pre- and post-normalization data, plotting the five and fifteen-thousand edge networks (Figure 5A, 5B). Rather than forming a handful of connected components centered around hub genes, RPCO-normalized data formed up to 2,073 discrete clusters for the fifteen-thousand edge network (Figure 5A). On the other hand, pre-normalization DepMap data represented nearly an order of magnitude fewer clusters for the fifteen-thousand edge network, 290, with the majority of edges instead concentrated into a single connected component with many mitochondrial-associated edges (Figure 5B). Comparing the number of genes represented across networks further illustrates that RPCO normalization uncovers relationships previously masked by mitochondrial-associated signal, with 10,493 more genes in the fifteen thousand-edge RPCO network than the corresponding pre-normalization network (Figure 5C).

**Figure 5:**
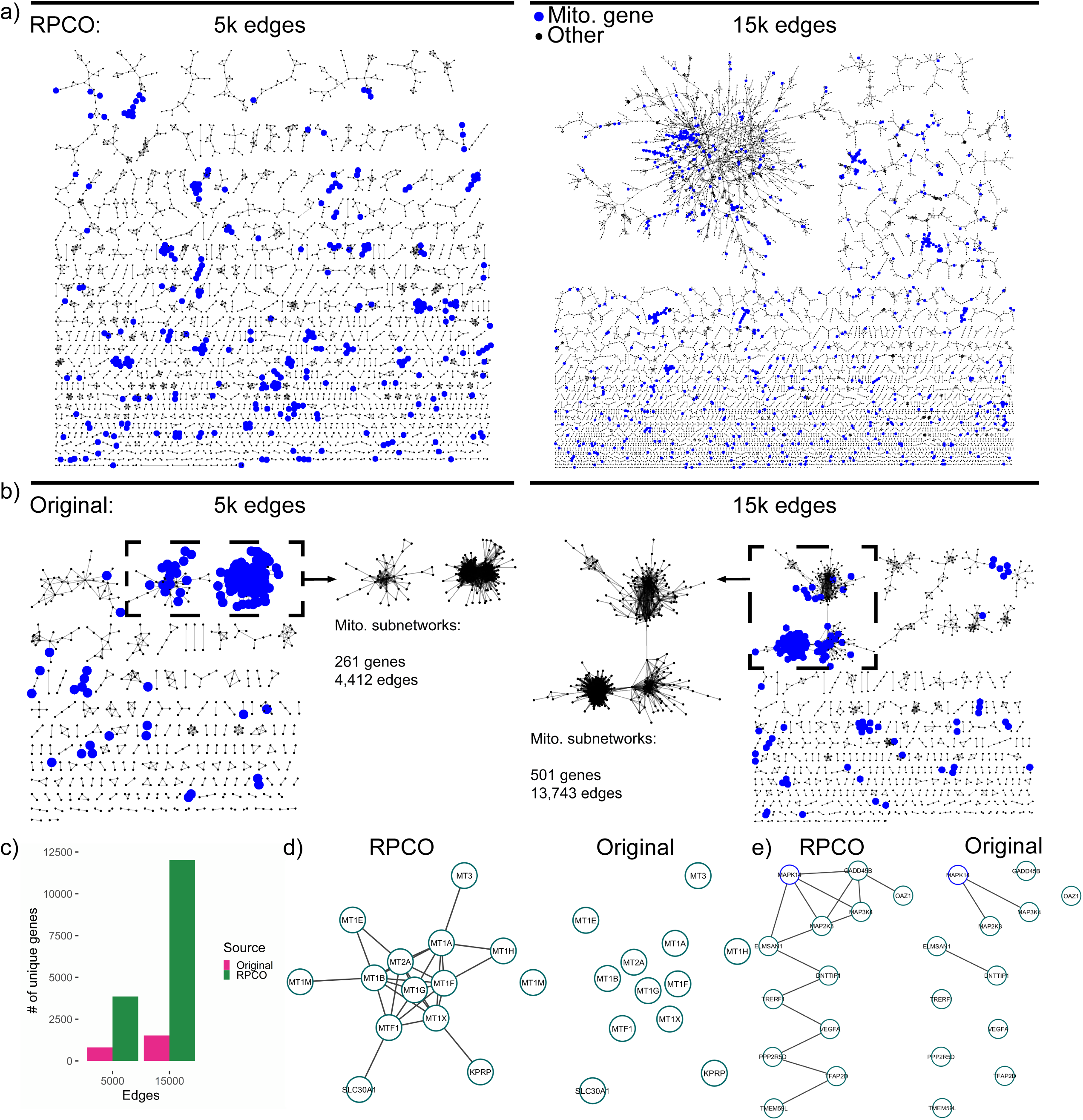
Network analysis of RPCO-normalized and original DepMap data. **a**, Top-ranked edges between genes from RPCO data laid out with the yFiles organic layout algorithm in Cytoscape (Shannon, et al., 2003), with mitochondrial-associated genes highlighted in blue. (Left) The top 5,000 edges for *n* = 3,850 genes. (Right) The top 15,000 edges for *n* = 12,017 genes. **b**, Top-ranked edges based on Pearson correlations between genes from original DepMap data laid out with the yFiles organic layout algorithm in Cytoscape (Shannon, et al., 2003), with mitochondrial-associated genes highlighted in blue. The largest connected components of the networks are inset and represent many mitochondrial-associated genes. (Left) The top 5,000 edges for *n* = 810 genes. (Right) The top 15,000 edges for *n* = 1,524 genes. **c**, The number of genes represented in above RPCO and original DepMap networks. **d**, Cluster derived from the 15,000 edge RPCO network representing metal homeostasis genes. (Left) Edges present in 15,000 edge RPCO network. (Right) Edges present in 15,000 edge original DepMap network. **e**, MAPK14-centric cluster derived from the 15,000 edge RPCO network. (Left) Edges present in 15,000 edge RPCO network. (Right) Edges present in 15,000 edge original DepMap network.

An investigation of clusters derived from RPCO-normalized data which lack signal in the original DepMap reveals potentially novel functions for the genes KPRP, DNTTIP1, TMEM59L and ELMSAN1. Twelve out of thirteen of a cluster of genes with a mean z-score of 43.8 in RPCO-normalized data, compared to a z-score of 1.9 in the original DepMap, are enriched for GO terms related to metal homeostasis (Figure 5D). The remaining gene, KPRP, is mostly uncharacterized and is not annotated to any GO biological process term. Therefore, we hypothesize that KPRP is also involved in metal homeostasis, perhaps working in conjunction with its nearest neighbor MT1X. A separate cluster of twelve genes, with a z-score of 30 in RPCO-normalized data compared to a z-score of 1.5 in the original DepMap, is enriched for MAP kinase signaling-related genes such as MAPK14 (Figure 5E). Intriguingly, while the gene ELMSAN1 (since renamed to MIDEAS) is known to be involved with histone deacetylation but little else, it is connected to both MAPK14 and MAP2K3. Through these connections, the similarly-uncharacterized genes DNTTIP1 and TMEM59L are associated with this cluster as well, indicating a potential connection between ELMSAN1, DNTTIP1, TMEM59L and MAPK14 activity.

### Onion-normalization enhanced signals in gene-expression data

To explore the generalizability of our onion normalization methods to other genome-scale datasets, we applied onion normalization to a single-cell gene expression dataset^31^ generated from healthy Peripheral blood mononuclear cells (PBMCs) using Chromium scRNA-seq technology and Cell Ranger (Data ref: (10xgenomics, 2019)). The pre-processed data contains log-normalized expression readouts for 12,410 genes across 1195 cells. A FLEX PR curve from the un-normalized data benchmarked against the CORUM protein standard shows the detection of 2,000 true positive gene pairs at a precision threshold of 0.8 (Figure S17A). However, the corresponding diversity plot shows that the majority of the strong performance (high precision) indicated by the PR curve comes from the cytoplasmic ribosome complex. PR-curves from RPCA- and RPCO-normalized data outperform un-normalized data by increasing the number of true positive (TP) gene pairs from 2,000 to 5,000 at a precision threshold of 0.8 (Figure S17B left). Moreover, PR curves without ribosomal gene pairs in the evaluation reveal that the normalized data performs better than the un-normalized data (Figure S17B right). For example, at a precision threshold of 0.2 the un-normalized data has around 50 TP gene pairs, whereas RPCO-normalization has 100. This suggests that the normalization process enhances signals in gene pairs within non-ribosomal complexes. For example, a closer look at the per-complex AUPRC values reveals that RPCO-normalization increased AUPRC for the Ferritin complex from 0.036 to 0.25 and for the Cofilin-actin-CAP1 complex from 0.028 to 0.146 (data not shown). This indicates that an optimized Onion-normalization method can be used to generally boost signals in gene expression data as well as CRISPR screen data.

## Discussion

In this study, we explored the use of unsupervised dimensionality techniques to identify functional relationships between genes within whole-genome CRISPR screening data and proposed a novel method called “onion” normalization for integrating signal between different “layers” of normalized data. While deep learning with autoencoders efficiently removed unwanted mitochondrial signal from the DepMap, this performance came at the expense of signal within smaller, non-mitochondrial complexes. Onion normalization rescued this poor performance for small complexes while still reducing mitochondrial signal and outperformed all proposed and state-of-the-art normalization methods when paired with robust principal component analysis (RPCO).

Co-essentiality maps derived from RPCO-normalized data show an unprecedented ability to recover signal from most of the genome when contrasted against the un-normalized DepMap and previous DepMap-derived co-essentiality maps. The fifteen-thousand edge RPCO network, constructed in a completely unsupervised way by measuring Pearson correlations above a given threshold, contained a total of ∼12k genes with an average of 2.5 neighbors per gene. The same approach applied to the original DepMap captured only ∼1,500 genes with an average of 19.7 neighbors per gene, likely due to the dominance of mitochondrial-associated hub genes within the network. Previous co-essentiality maps constructed from the DepMap either filtered out the majority of the genome or initialized the network structure based on a set of pre-existing clusters^6,7^ (Wainberg, et al., 2021; Kim, et al., 2019), techniques ill-suited for mapping the functions of understudied genes. RPCO-normalization overcomes these limitations and allows us to ascribe putative functions to previously weakly-connected genes.

Our exploration provides a compendium of resources for studying functional relationships within the DepMap at an improved resolution, including a novel co-essentiality map and the onion normalization method. While our results show a strong performance benefit for robust principal component analysis, future work could investigate both deep learning approaches for normalizing the DepMap and onion normalization applied to different input normalization approaches. Perhaps other deep-learning approaches that learn meaningful latent spaces, such as variational autoencoders (Kingma & Welling, 2013), could better learn and remove mitochondrial signal without reducing signal within mitochondrial-associated complexes. As the key technical limitation of onion normalization is its high memory cost, which scales with the number of layers, future work could also investigate the choice of optimal hyperparameters across different layers of normalized data. Additionally, onion normalization is a general framework that our initial analyses suggest may be applicable to other types of genomic data such as bulk and single-cell RNA-seq.

## Methods

### Principal component analysis normalization

We applied the R function prcomp (an SVD-based R implementation of PCA) to the original DepMap 20Q2 data (Data ref: (Broad, 2020)) with scale and center parameters set to true and generated corresponding principal component (PC) outputs. Prior to that, NA values were replaced with gene-wise mean CERES scores in the downloaded DepMap data (Achilles_gene_effect.csv). The rotation variable of the PCA output corresponds to loadings of the principal components. Multiplying DepMap CERES scores with the complete rotation matrix transforms the data to a coordinate space defined by the principal components. Multiplying this resulting matrix with the transpose of the loadings matrix re-transforms data into the original coordinate space. In our method, the original data matrix *M_r_*_×*c*_ multiplied by only a subset of the principal component loadings matrix (*L_c_*_×*c*_) and its transpose. This creates a ‘PCA-reconstructed’ version of the original data matrix from the low dimensional signal-space defined by that particular subset of principal components (Equation 1.1). The *n*-PC PCA-reconstruction of the original data is thus generated using the first *n* columns of the rotation matrix. Subtracting the PCA-reconstructed matrix from the original data matrix generates the *n*-PC removed PCA-normalized version of the data (Equation 1.2).

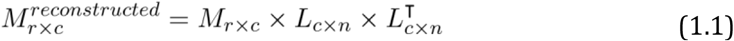

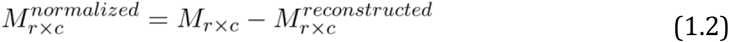

### Robust principal component analysis normalization

Robust principal component analysis (RPCA) decomposes matrix *X_r_*_×*c*_ into low-rank, *L_r_*_×*c*_, and sparse, *S_r_*_×*c*_, component matrices so that they satisfy Equation 1.3 (Candès, Li, Ma, & Wright, 2011). RPCA is an unsupervised method, designed to optimize the values of *L* and *S* to minimize Equation 1.4, where ||*L*||_∗_ is the nuclear-norm of *L* and ||*S*||1_∗_ is the *l*_1_-norm of *S*. *λ* is a hyperparameter whose suggested value is 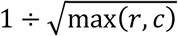

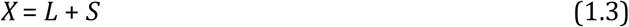

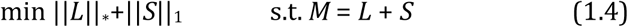

For this work, we applied the rpca R-package (an R implementation of RPCA) (Sykulski, 2015) to the original DepMap data (Data ref: (Broad, 2020)). As a pre-processing step, NA values replaced with gene-wise mean CERES scores in the downloaded DepMap data (Achilles_gene_effect.csv). Variables *S* and *L* in rpca output are the RPCA-normalized data and RPCA-reconstructed data, respectively.

### Autoencoder normalization

The 20Q2 DepMap data, Achilles_gene_effect.csv (Data ref: (Broad, 2020)), was processed in the following way to prepare data for fitting with an autoencoder model. First, NA values were replaced with gene-wise mean CERES scores. Second, the dataset was row-standardized. Third, the 0.12% of resulting z-scores below −4 or above 4 were clipped to −4 or 4, respectively. Lastly, the entire dataset was min-max scaled to fall between −1 and 1.

A deep convolutional autoencoder was then trained on the DepMap for 1 epoch and a latent space size of *LS* = 1, 2, 3, 4, 5 or 10. The encoder architecture consisted of a 1D convolutional layer converting from 1 channel into 10 with a subsequent 1D max pooling layer, another 1D convolutional layer converting from 10 channels into 20 with a subsequent 1D max pooling layer, and flattening followed by a linear layer with size equal to the chosen latent space. The decoder architecture consisted of inverse operations with max unpooling, transposed convolutional layers and a final linear layer to reshape output into the original input size. All convolutional kernel sizes were set to 3 and all pooling kernel sizes were set to 2.

### Onion normalization

The onion normalization method combines signals from different normalized data (that we refer to as ‘layers’) generated by dialing parameter values of a normalization method. It has three components - normalizing gene effect scores with a dimensionality reduction method, creating similarity-networks from normalized data, and finally integrating the similarity-networks into a single network.

Any effective dimensionality reduction method can be employed in the normalization step. The layers to be fused are produced from the same data normalized by varying a parameter of the normalization method. We created such layers by applying PCA, RPCA or AE normalization methods as described in their respective sections. For example, we created six AE-normalized layers using AE-normalization with latent space sizes of 1, 2, 3, 4, 5, and 10. Similarly, we removed the first *n* principal components (for *n* = 1, 3, 5, 7, 9, 11, 13, 15, 17, or 19) and generated ten PCA-normalized layers. For RPCA-normalization, we regulated *λ* applying the formula *f*÷ 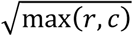 for *f* = 0.7, 0.8, 0.9, 1, 1.1, 1.2, 1.3 and generated seven RPCA-normalized layers for integration. From each normalized layer, we created a gene-level similarity network by computing Pearson correlation coefficients among the gene profiles.

For the network integration module of the Onion method, we selected the Similarity Network Fusion (SNF) approach developed by Wang et. al. as it outperformed baseline integration techniques we explored (Figure S6, Figure S7, Figure S8) (Wang, et al., 2014). SNF is a network fusion technique based on multiview learning that enhances or diminishes network edge weights by considering the neighborhood and sparsity information of the individual networks. We converted Pearson correlation coefficients to distance metrics by subtracting them from 1 before applying a scaled exponential similarity kernel (the affinityMatrix function in the SNF package) to generate an affinity score matrix. These affinity matrices generated from each layer of normalized data are then integrated into one network with the SNF package.

SNF has three relevant hyperparameters. The first parameter, *σ*, is a standard deviation regulator of the exponential similarity kernel and is used to create the affinity matrices. Another hyperparameter, *k*, regulates the number of neighboring vertices to be considered during calculating edge weights in the integrated network and is used both in the affinity matrix creation and the final integration stages. A third hyperparameter controls the number of iterations in the integration stage. We dialed *σ* = 0.1, 0.3, 0.5, 0.7 and *k* = 3, 5, 10, 20 in integrating AE, PCA and RPCA normalized layers and settled on *σ* = 0.3, *k* = 5 for PCO and *σ* = 0.5, *k* = 5 for RPCO and AEO, based on how much diversity it can introduce during the evaluation process (Figure S5). We set the number of iterations to 10 for all methods. While integrating AE-normalized layers, we also included the similarity network generated from the un-normalized data as a layer, fusing a total of seven layers.

### Functional evaluations

To evaluate normalization methods we used the CRISPR screen benchmarking package FLEX and the CORUM (Giurgiu, et al., 2019) protein complex database as FLEX’s gold standard to benchmark against (Rahman, et al., 2021). FLEX’s evaluation is based on the idea that gene-level similarity scores, calculated from gene knock-out profiles, connotes functional similarity among genes and a higher similarity score between two genes implies membership in the same protein complex. FLEX orders gene pairs from high to low similarity scores and evaluates complex membership predictions at different precision points against the CORUM standard. The precision-recall (PR) curve from FLEX depicts how many true positive (TP) gene pairs are both strongly correlated within the data and members of the same CORUM protein complex. This visualization is augmented by diversity plots, which illustrate specific complexes that contain most true-positive gene pairs at various precision points. A visually larger area denotes more TP contribution from a complex. Another evaluation metric in FLEX is the per-complex area under the PR curve (AUPRC) value. In calculating AUPRC for a complex, gene pairs belonging to that complex are considered as positive examples whereas gene pairs from other complexes are set as negative examples. A higher per-complex AUPRC indicates more gene pairs associated with that complex have been identified based on their similarity scores. Conversely, a lower per-complex AUPRC means that scores for the within-complex genes are poorly correlated compared to between-complex gene pairs.

FLEX also facilitates removing specific gene pairs from the evaluation process. To evaluate the influence of mitochondrial complexes in the DepMap data, we compiled 1,266 mitochondrial genes from three sources to remove from our analysis. A total of 1,136 genes were collected from the Human MitoCarta3.0, an inventory of human mitochondrial proteins and pathways by the Broad Institute (Rath, et al., 2021). All genes from the KEGG OXIDATIVE PHOSPHORYLATION and the REACTOME RESPIRATORY ELECTRON TRANSPORT pathways were included in the list. 436 genes were also assembled by an expert based on information from pathways and CORUM complexes, and the union of these lists formed a reference list of mitochondrial-associated genes. To modify FLEX analyses according to this list and better examine non-mitochondrial signal within the DepMap, gene pairs were excluded from FLEX analyses for pairs where both genes are contained in the mitochondrial gene list. Gene pairs that contain only one or no mitochondrial genes are not removed.

### Network analysis

Networks were constructed from the original 20Q2 DepMap and the RPCO-normalized 20Q2 DepMap datasets by taking the top five, ten, or fifteen-thousand edges based on the strength of Pearson correlations across each respective dataset. Network layouts were performed with the yFiles organic layout algorithm in Cytoscape version 3.7.2 (Shannon, et al., 2003). All connected components within each network were treated as separate clusters and analyzed for enrichment. Enrichments tests were performed with hypergeometric tests using the clusterProfiler R package version 3.16.1 (Wu, et al., 2021) against human Gene Ontology-biological process and MSigDB C2 curated pathway annotations and a background set of all genes in the given network at a Benjamini-Hochberg FDR of 0.2.

### Analysis on gene-expression data

The scRNA-seq gene expression dataset (5k_pbmc_v3_filtered_feature_bc_matrix.tar) was downloaded from 10xGENOMICS (10xgenomics, 2019) and was generated from Peripheral blood mononuclear cells (PBMCs) using Chromium and Cell Ranger. We applied the Seurat R package to filter the dataset and removed genes for which the number of cells with non-zero values is smaller than or equal to 50. We also filtered out cells for which the number of unique genes detected in each cell is smaller than or equal to 100 and greater than or equal to 4500. Furthermore, we only included cells for which the percentage of reads that map to the mitochondrial genome is lower than 7. The final matrix contains 12410 genes and 1195 cells, around 20% of which is non-zero. We log-normalized the data using Seurat function NormalizeData with default parameters.

We applied RPCA to the pre-processed scRNA-seq data and generated seven RPCA-normalized layers by setting hyperparameter lambda to f÷√max(r,c), where r = 12000, c=1200, and f = 0.7, 0.8, 0.9, 1, 1.1, 1.2, 1.3. Seven gene-gene similarity networks were generated from the normalized data using Pearson Correlation Coefficients as the similarity metric. The networks were integrated by taking the maximum weight for each gene-pair across the seven networks. To demonstrate the dominance of cytoplasmic ribosomal gene pairs in the analysis results (Figure S17B right), we removed 81 Ribosome (cytoplasmic) complex-associated genes during the FLEX evaluation process.

## Acknowledgements

This research was funded by grants from the National Science Foundation (MCB 1818293), the National Institutes of Health (R01HG005084, R01HG005853). H.N.W was partially supported by the Doctoral Dissertation Fellowship from the U. of Minnesota and a National Institutes of Health (NIH) Biotechnology Training Grant (T32GM008347). M.B. was partially supported by the German Research Foundation DFG (Bi 2086/1–1).

## Author contributions

Conceptualization: AZH, HNW, CLM; Software: HNW, AZH, MR;

Formal Analysis: HNW, AZH;

Investigation: HNW, AZH, MB, MR, YL, CLM; Resources: MB, MR, YL;

Data Curation: HNW, AZH;

Writing – Original Draft: HNW, AZH;

Writing – Review & Editing: HNW, AZH, CLM, MB, MR, YL; Visualization: HNW, AZH;

Supervision: CLM;

Project Administration: CLM; Funding Acquisition: CLM;

## Data availability

The datasets and computer code produced in this study are available in the following databases:

- Codes to reproduce the main figures: GitHub (https://github.com/ArshiaZHassan/ONION_git)
- Codes for autoencoder-normalization: GitHub (https://github.com/csbio/ae-norm)
- Data to reproduce the main figures and associated outputs: Zenodo https://doi.org/10.1101/2023.02.22.529573 (https://zenodo.org/record/7671685#.Y_gi9nbMK5c)

---

**Figure S1:**
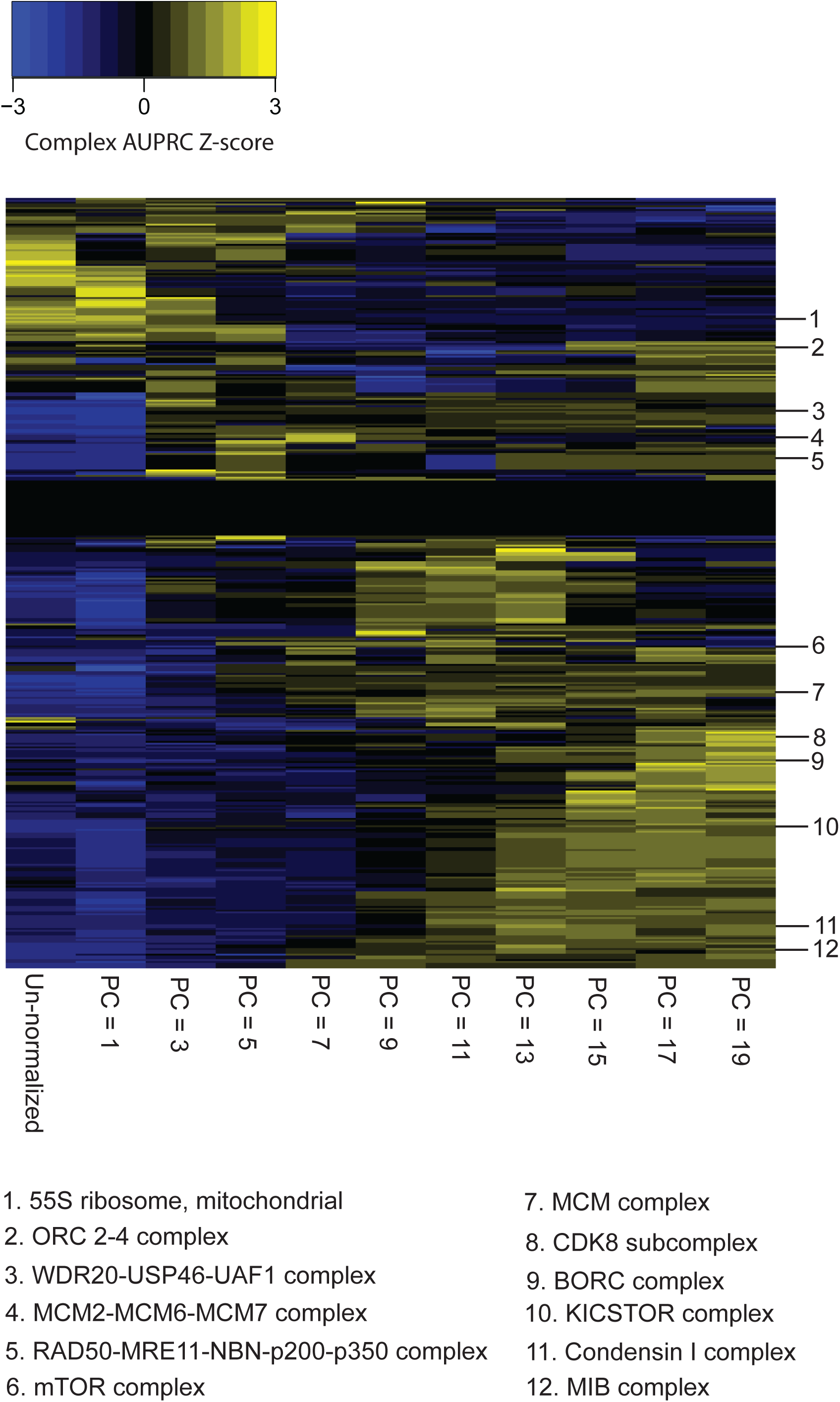
CORUM complex z-scores of AUPRC values for PCA normalization. Z-scores across rows for un-normalized DepMap data compared to DepMap data with the first 1, 3, 5, 7, 9, 11, 13, 15, 17, or 19 PCs removed.

**Figure S2:**
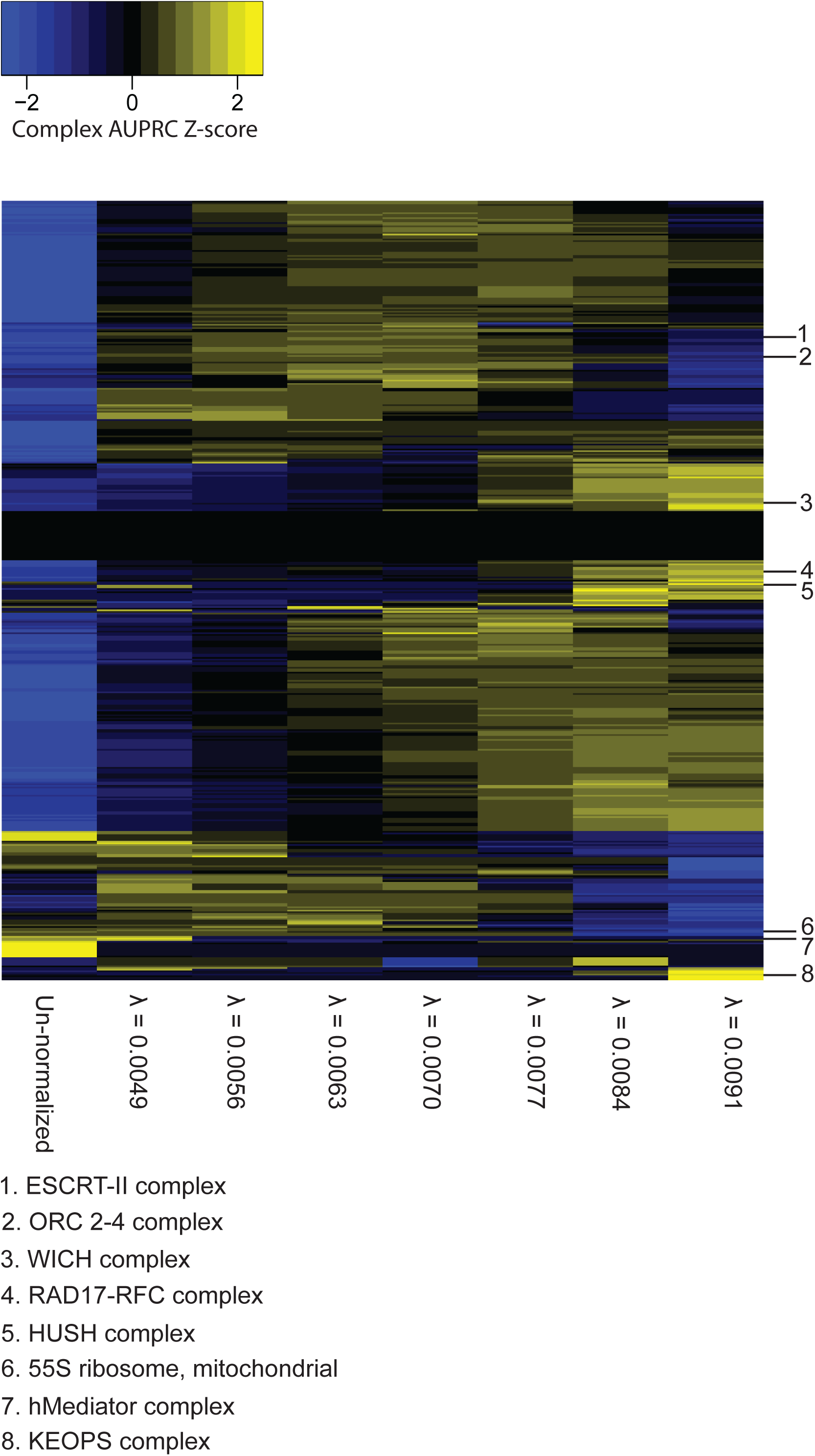
CORUM complex z-scores of AUPRC values for robust PCA normalization. Z-scores across rows for un-normalized DepMap data compared to robust PCA-normalized DepMap data for *λ* ~= 0.0049, 0.0056, 0.0063, 0.007,0.0077, 0.0084, 0.0091.

**Figure S3:**
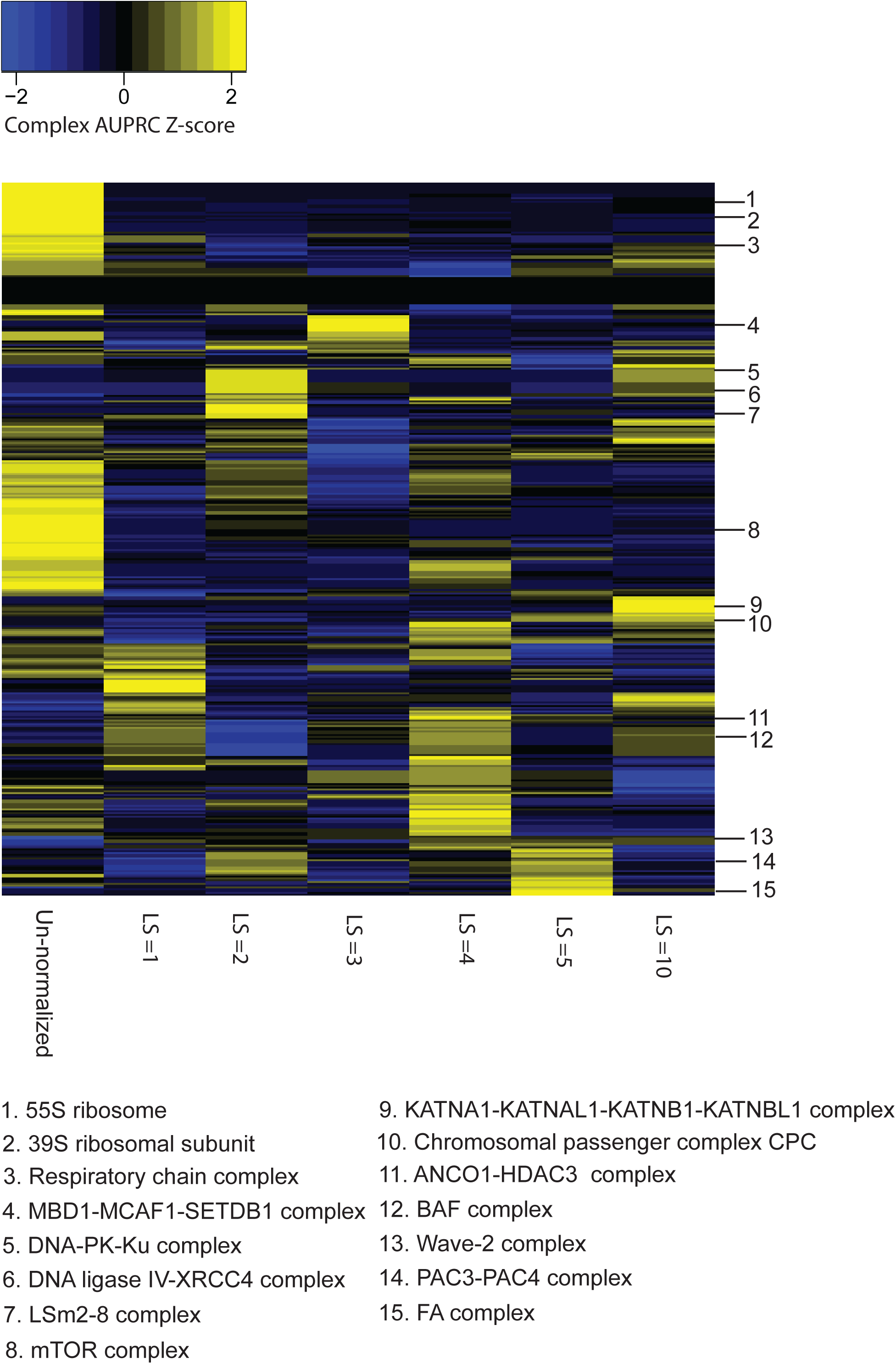
CORUM complex z-scores of AUPRC values for autoencoder normalization. Z-scores across rows for un-normalized DepMap data compared to autoencoder-normalized DepMap data with latent space size (LS) = 1, 2, 3, 4, 5, 10.

**Figure S4:**
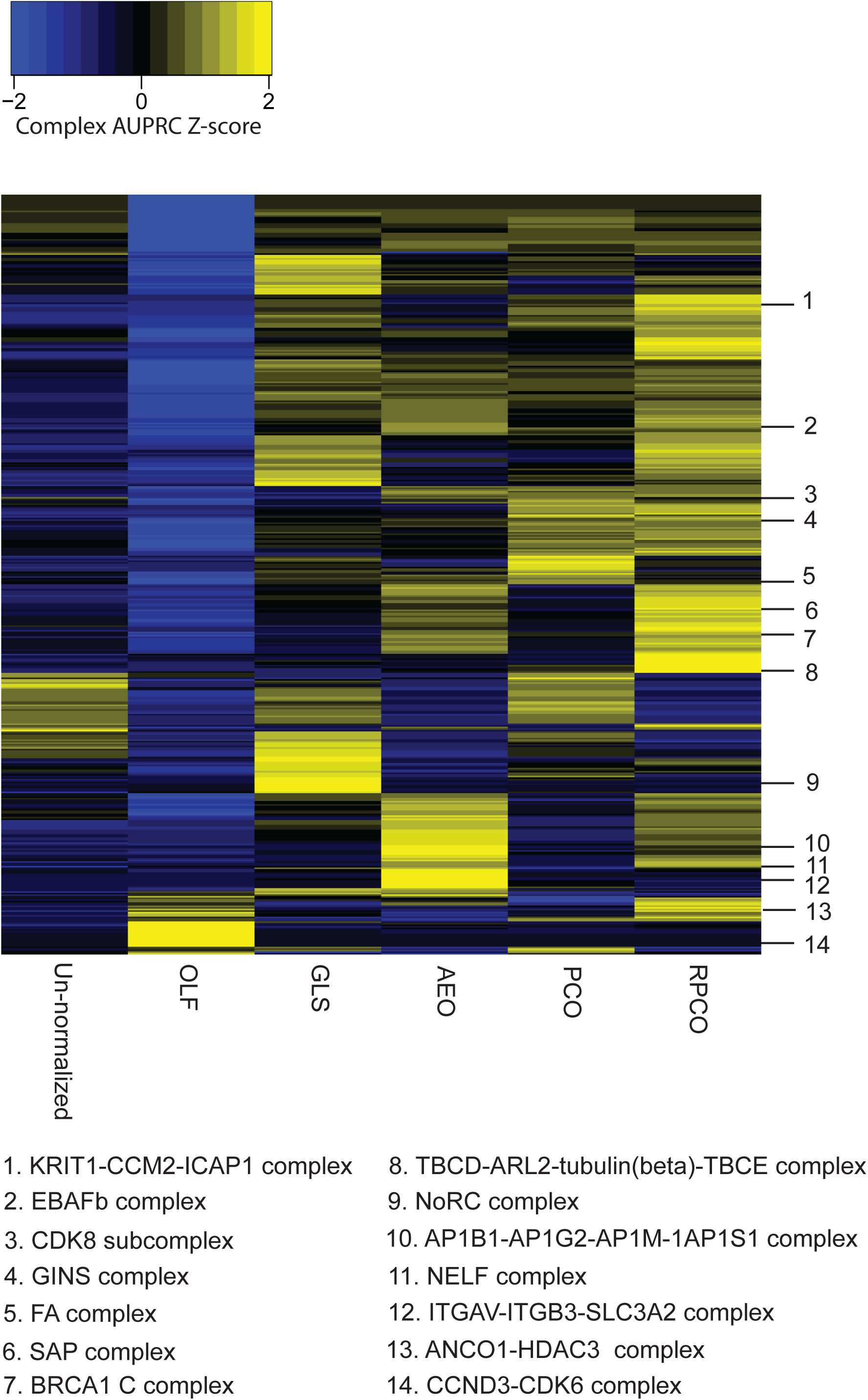
CORUM complex z-scores of AUPRC values for onion normalization. Z-scores across rows for un-normalized DepMap data compared to data from generalized least squares (GLS) normalization from Wainberg et al. (Wainberg, et al., 2021), olfactory receptor (OLF) normalization from Boyle et al. (Boyle, Pritchard, & Greenleaf, 2018), onion normalization with AE (AEO), onion normalization with PCA (PCO), and onion normalization with robust PCA (RPCO).

**Figure S5:**
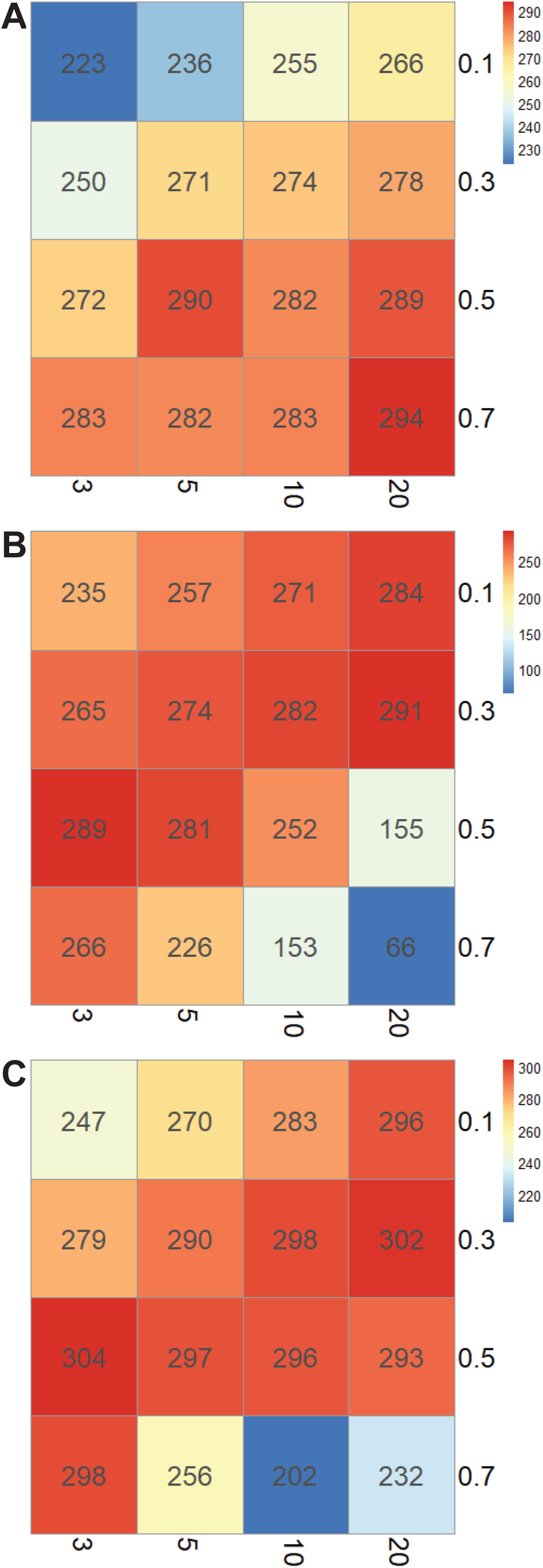
SNF hyperparameter exploration for onion normalization. The number of CORUM protein complexes which contribute any true positive pairs at a precision threshold of 0.5 are plotted for SNF parameters of *σ* = 0.1, 0.3, 0.5, 0.7 on the y-axis and *k* = 3, 5, 10, 20 on the x-axis. **A**, Autoencoder-normalized data as input to onion normalization. **B**, PCA-normalized data as input to onion normalization. **C**, Robust PCA-normalized data as input to onion normalization.

**Figure S6:**
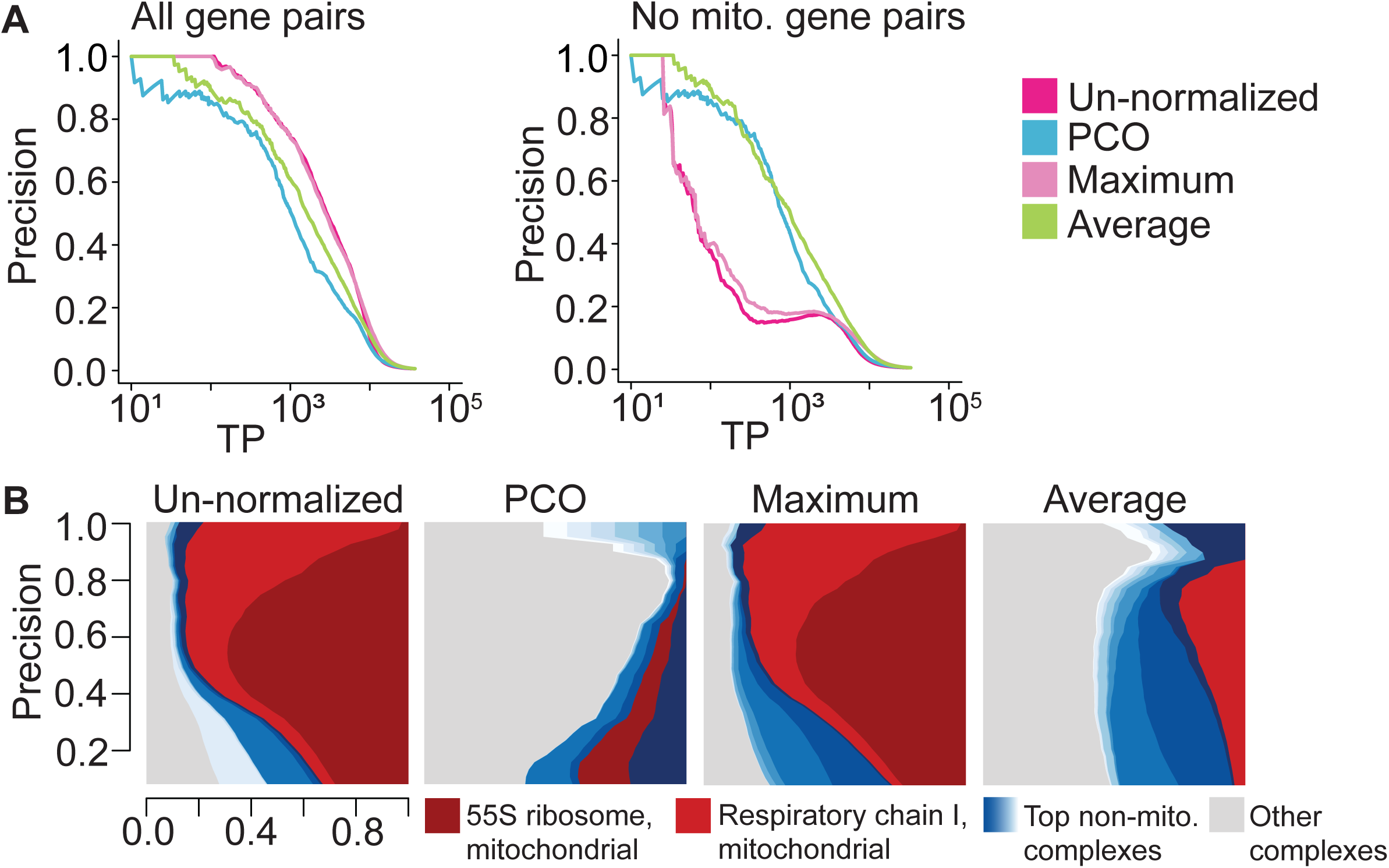
A, Comparison of Precision-recall (PR) performance of original DepMap data, SNF integrated PCA-normalized layers (PCO), average of PCA-normalized layers and maximum across PCA-normalized layers evaluated against CORUM complex standard. Layers are generated by removing first n principal components where n = 1, 3, 5, 7, 9, 11, 13, 15, 17, 19. **B,** Contribution diversity plots from original DepMap data, SNF integrated PCA-normalized layers (PCO), average of PCA-normalized layers and maximum across PCA-normalized layers evaluated against CORUM complex standard.

**Figure S7:**
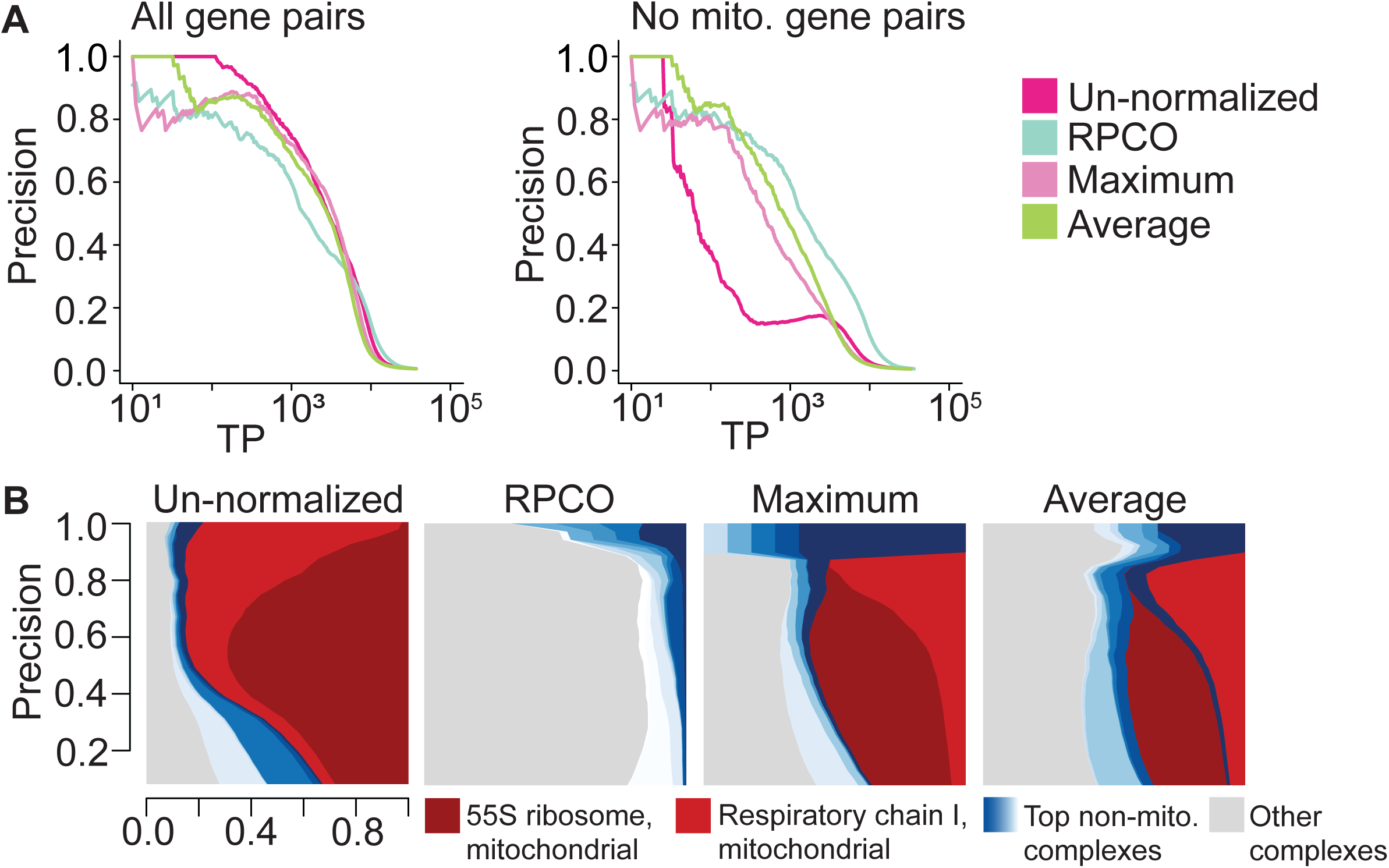
A, Comparison of Precision-recall (PR) performance of original DepMap data, SNF integrated RPCA-normalized layers (RPCO), average of RPCA-normalized layers and maximum across RPCA-normalized layers evaluated against CORUM complex standard. Layers are generated by setting RPCA hyperparameter *λ* to approximately 0.0049, 0.0056, 0.0063, 0.007, 0.0077, 0.0084, and 0.0091. **B,** Contribution diversity plots from original DepMap data, SNF integrated RPCA-normalized layers (RPCO), average of RPCA-normalized layers and maximum across RPCA-normalized layers evaluated against CORUM complex standard.

**Figure S8:**
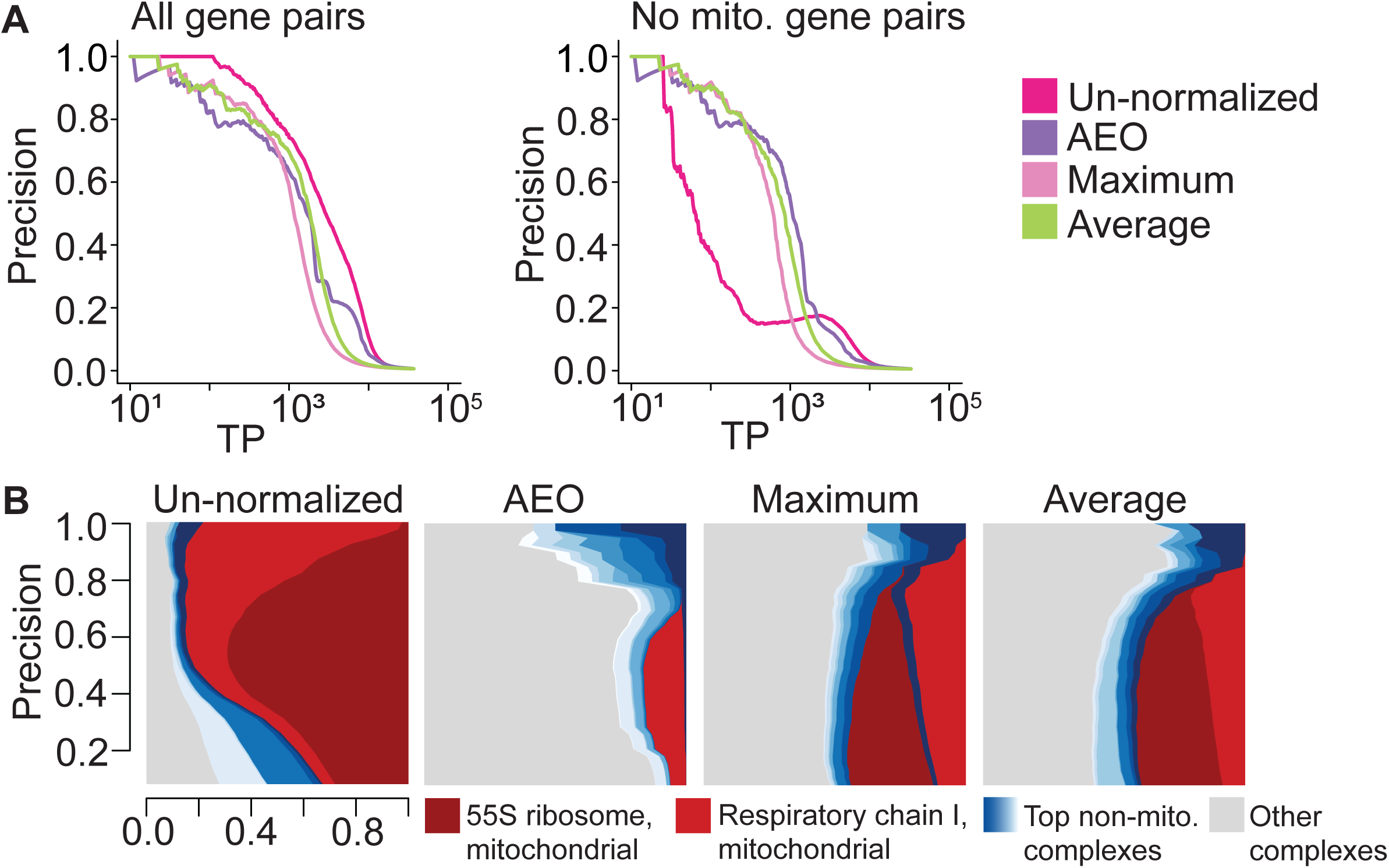
A, Comparison of Precision-recall (PR) performance of original DepMap data, SNF integrated AE-normalized layers (AEO), average of RPCA-normalized layers and maximum across RPCA-normalized layers evaluated against CORUM complex standard. Layers are generated by dialing auto-encoder latent space size to 1, 2, 3, 4, 5, and 10. **B,** Contribution diversity plots from original DepMap data, SNF integrated AE-normalized layers (AEO), average of RPCA-normalized layers and maximum across RPCA-normalized layers evaluated against CORUM complex standard.

**Figure S9:**
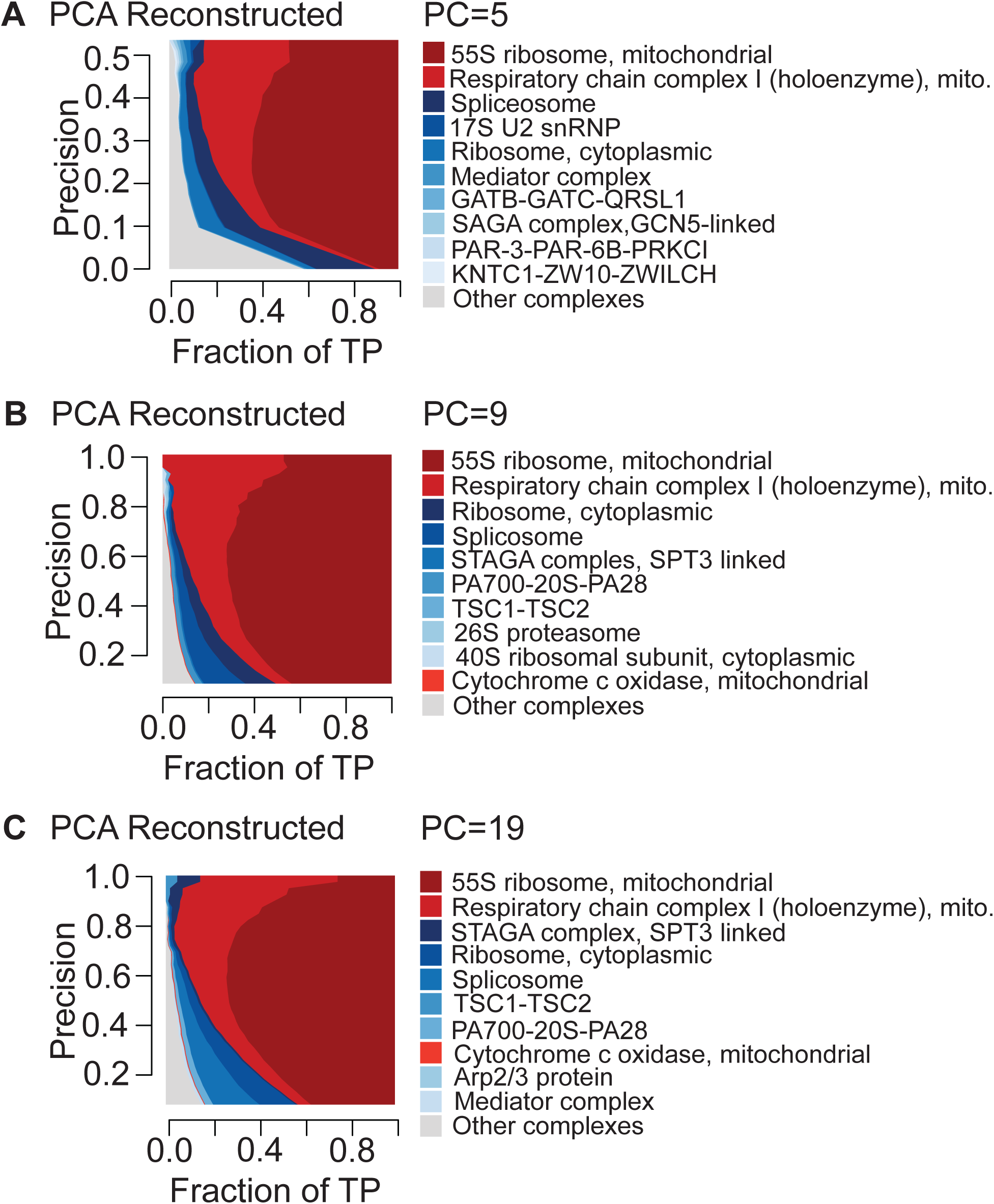
Contribution diversity plots depicting CORUM complex contributions in PCA-reconstructed data with first 5, 9 and 19 principal components removed.

**Figure S10:**
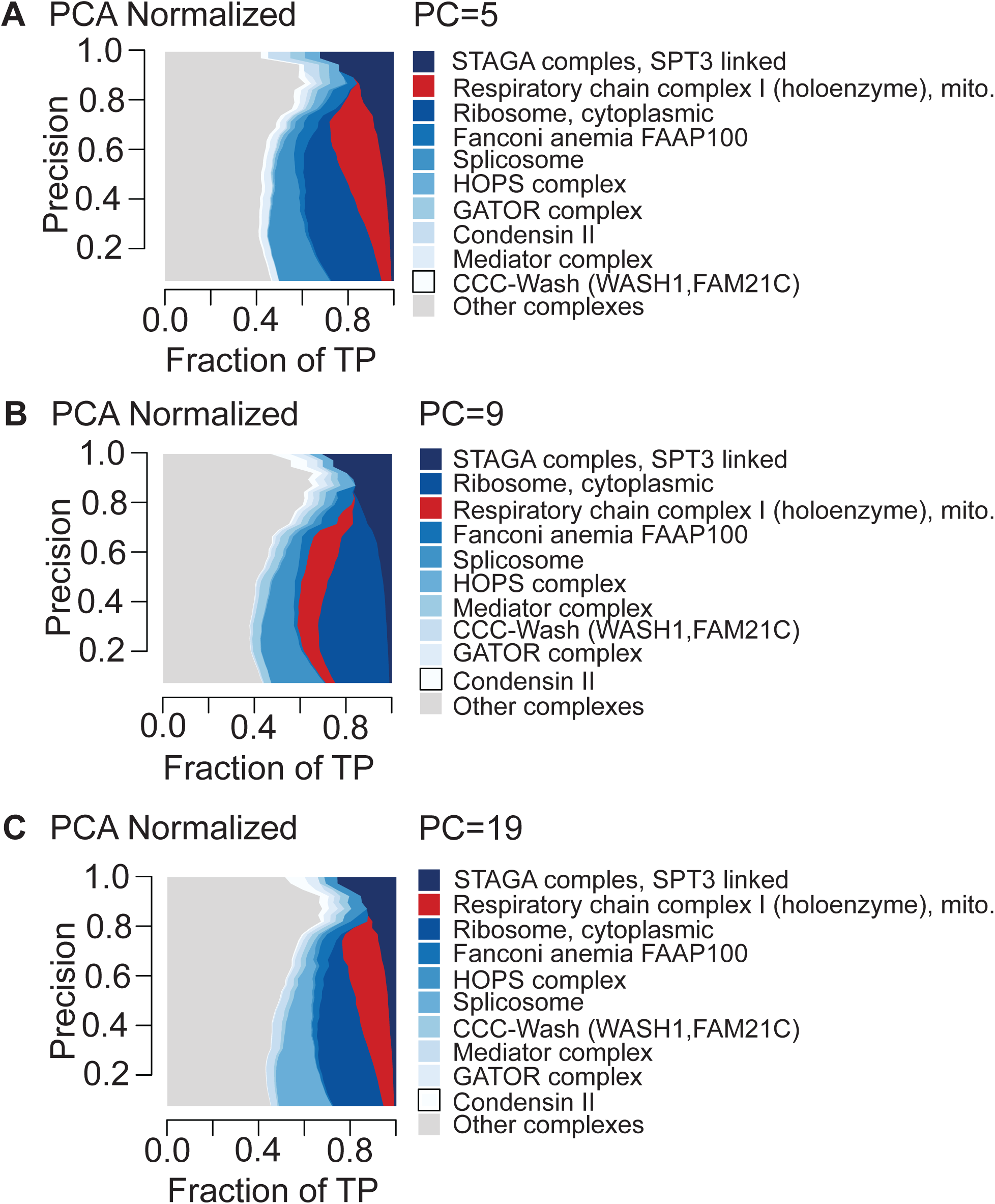
Contribution diversity plots depicting CORUM complex contributions in PCA-normalized data with first 5, 9 and 19 principal components removed.

**Figure S11:**
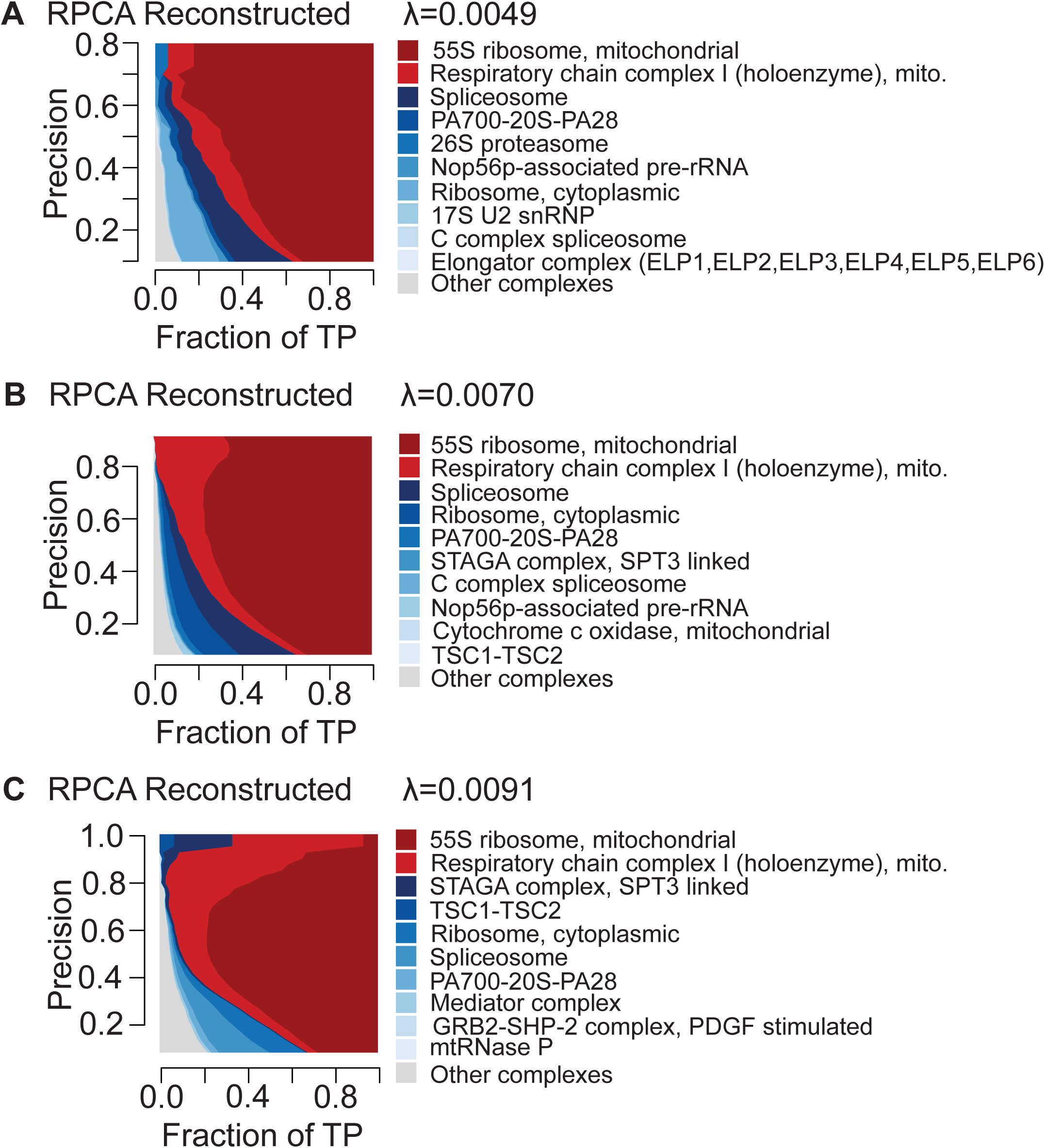
Contribution diversity plots illustrating complex contributions in RPCA-reconstructed data generated with hyperparameter *λ* set to 0.0049, 0.007 and 0.0091 evaluated against the CORUM complex standard.

**Figure S12:**
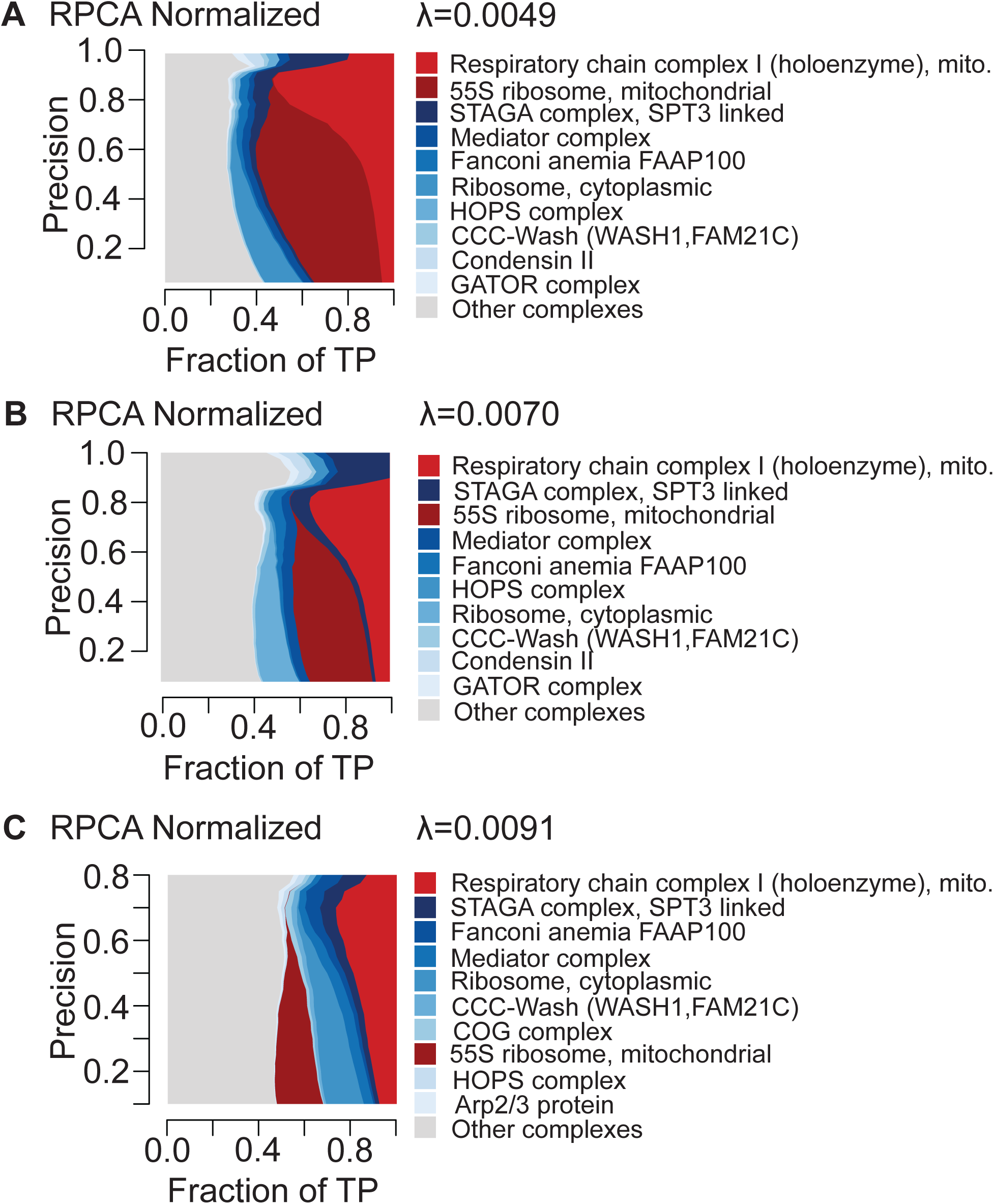
Contribution diversity plots illustrating complex contributions in RPCA-normalized data generated with hyperparameter *λ* set to 0.0049, 0.007 and 0.0091 evaluated against the CORUM complex standard.

**Figure S13:**
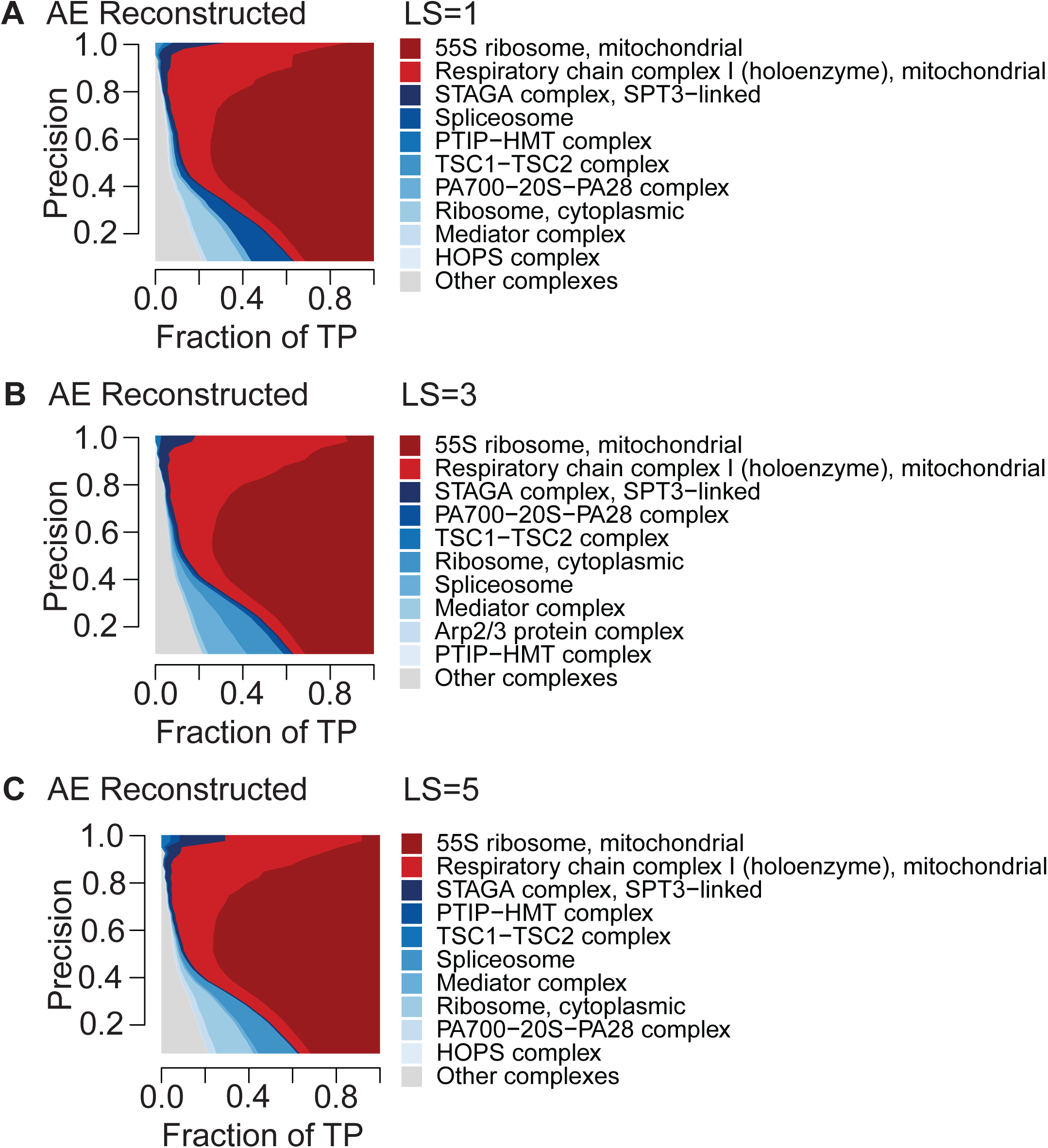
Contribution diversity plots depicting CORUM complex contributions from AE-reconstructed data generated with latent space sizes 1, 3 and 5 evaluated against CORUM complex standard.

**Figure S14:**
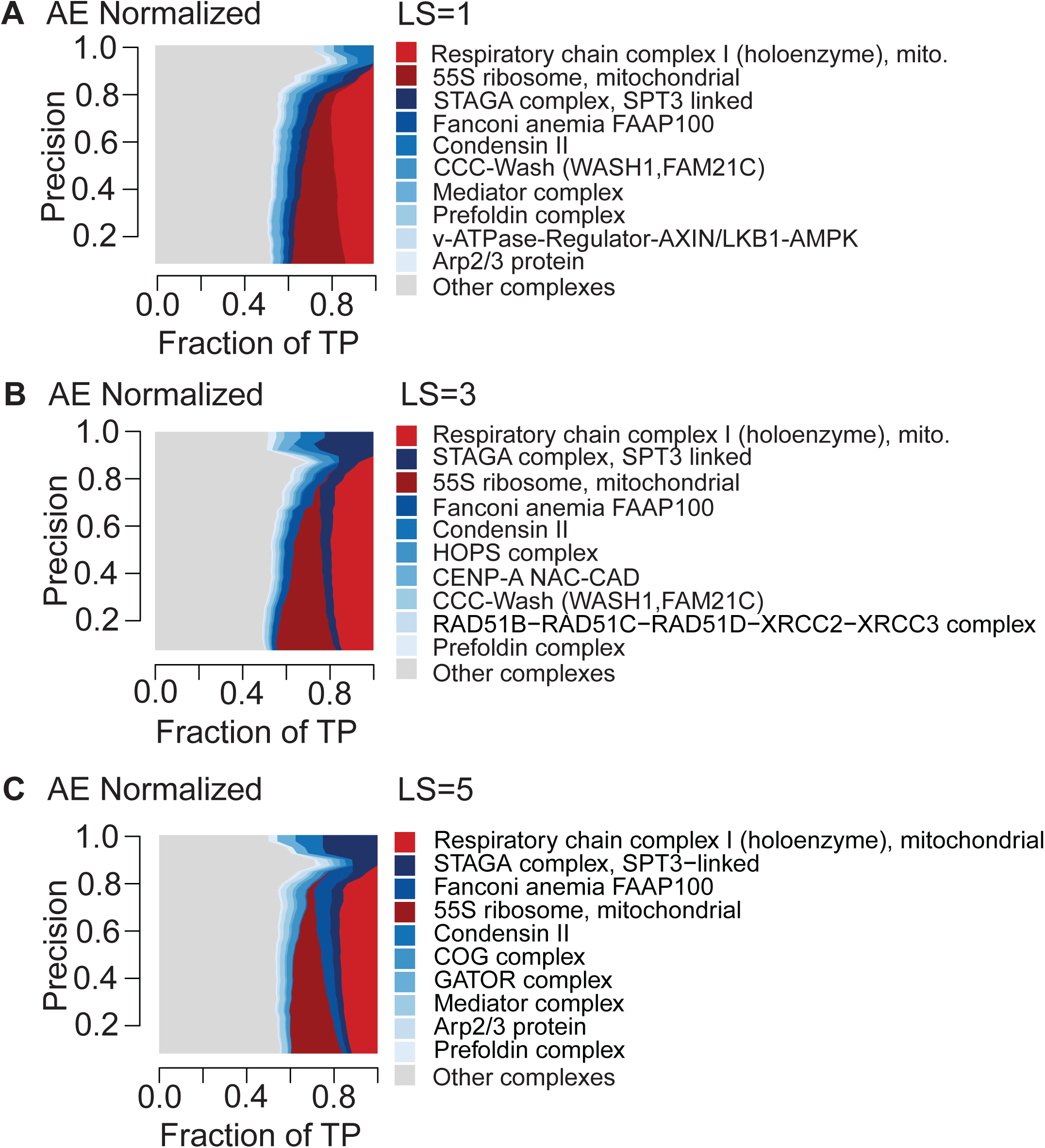
Contribution diversity plots depicting CORUM complex contributions from AE-normalized data generated with latent space sizes 1, 3 and 5 evaluated against CORUM complex standard.

**Figure S15:**
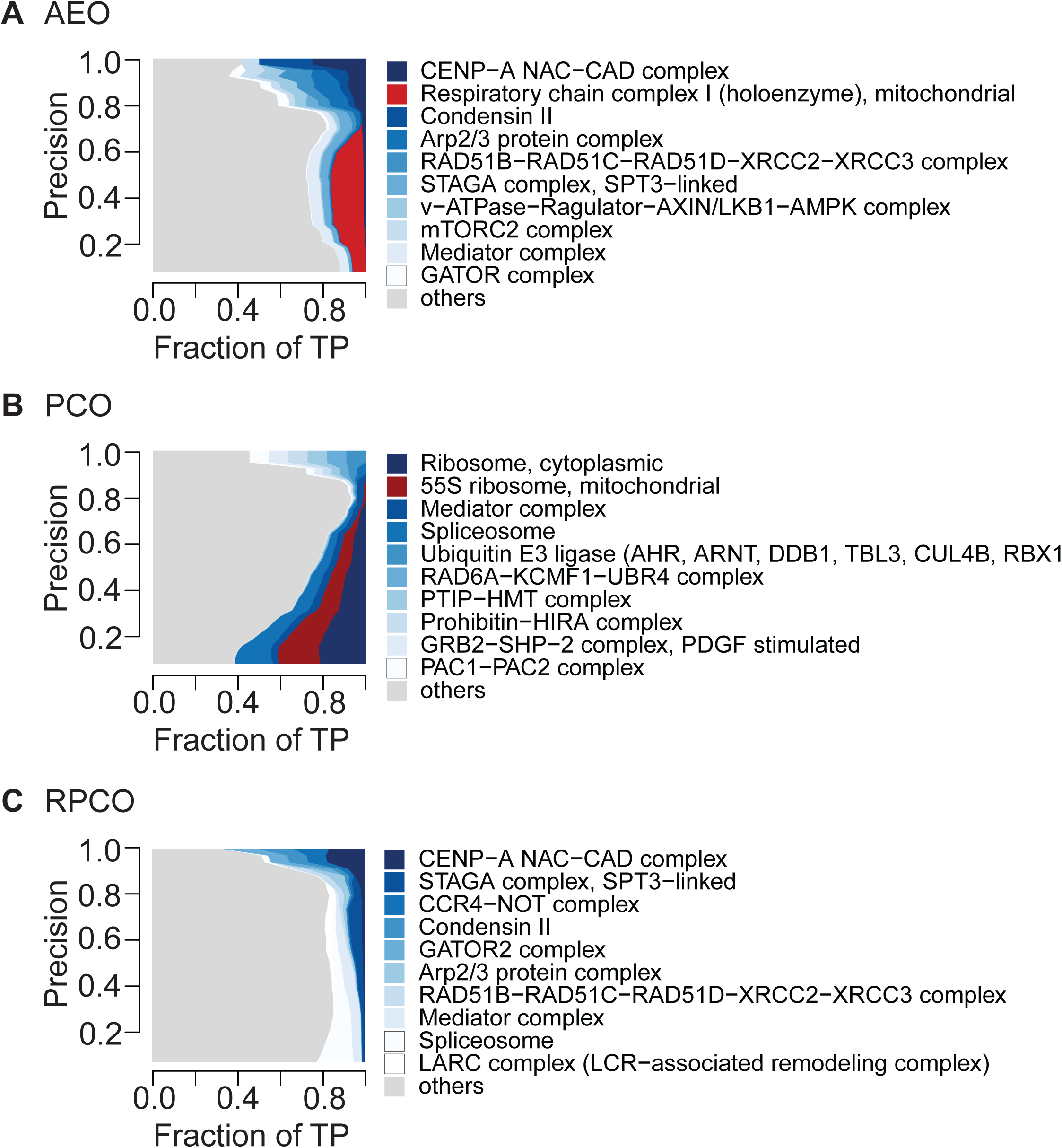
Contribution diversity of CORUM complexes in SNF integrated AE-normalized layers (AEO), SNF integrated PCA-normalized layers (PCO), and SNF integrated RPCA-normalized layers (RPCO).

**Figure S16:**
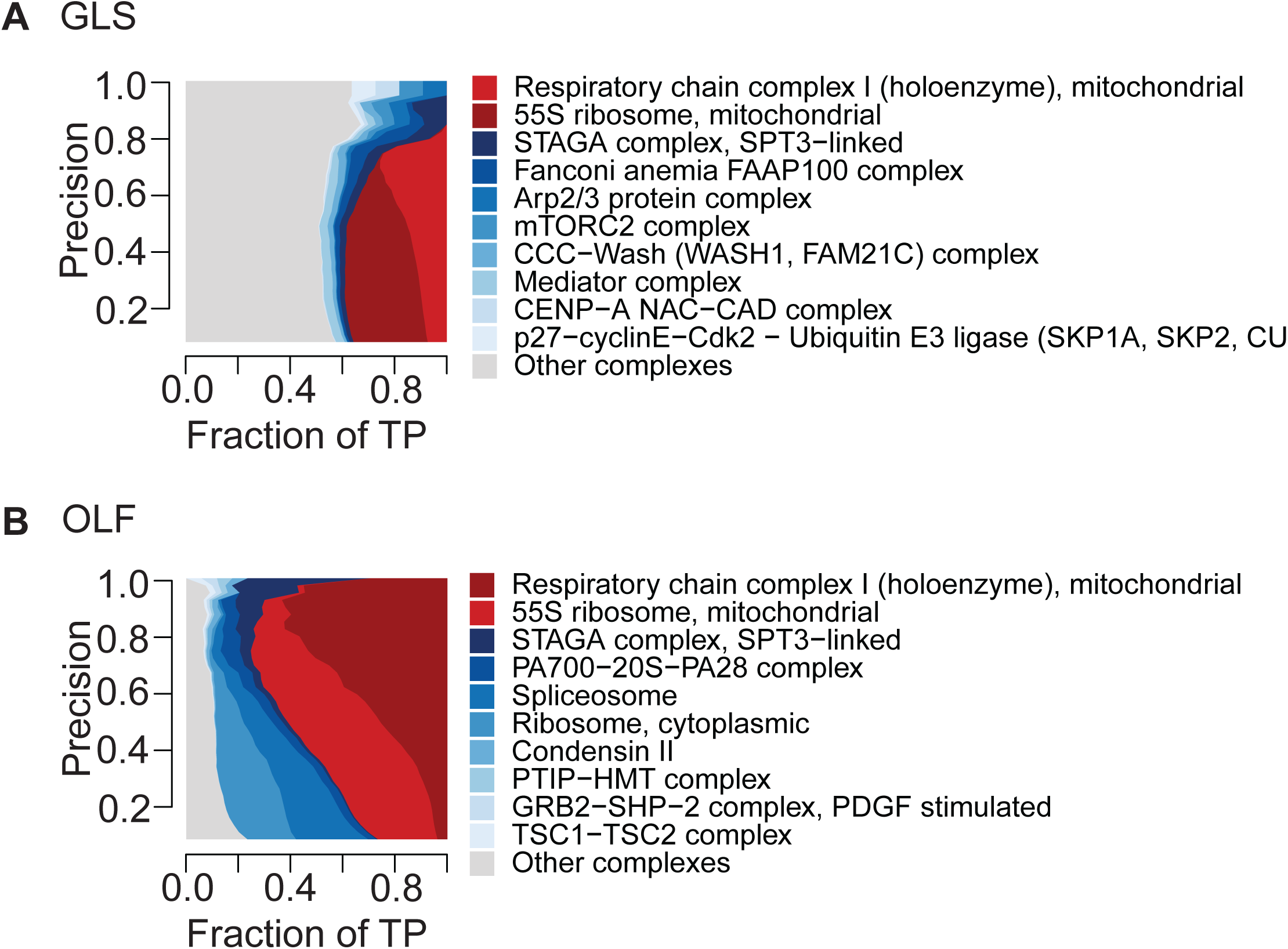
Contribution diversity of CORUM complexes in generalized least squares (GLS) normalization from Wainberg et al. (Wainberg, et al., 2021), olfactory receptor (OLF) normalization from Boyle et al. (Boyle, Pritchard, & Greenleaf, 2018)

**Figure S17:**
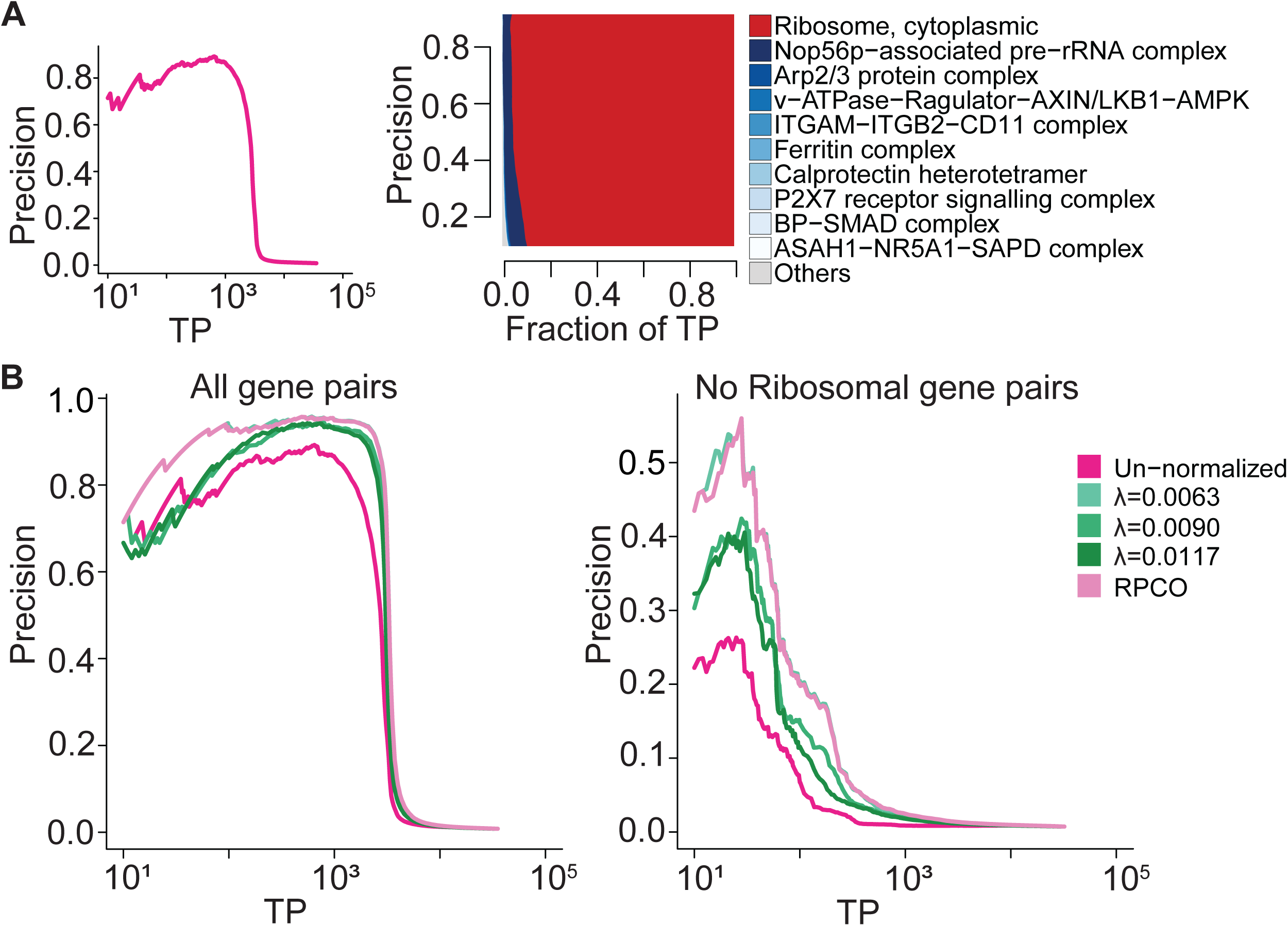
Onion-normalization applied to RNA-seq gene expression data with RPCA as the normalization technique and maximum-weight as integration method. **A,** (Left) Precision-recall (PR) performance of un-normalized gene expression data evaluated against CORUM protein complexes. The x-axis depicts the log-scaled number of true-positives (TPs). (Right) Contribution diversity of CORUM complexes in un-normalized gene expression data. The x-axis is the fraction of TP gene pairs against those pairs correctly predicted as functionally related at different precision levels in the y-axis. The legend shows the top ten contributing complexes. **B,** (Left) PR performance of RPCA-normalized data generated with λ set to 0.0063, 0.009, and 0.0117 as well as RPCO-normalized data evaluated against CORUM protein complexes. (Right) PR performance of RPCA-normalized data generated with λ set to 0.0063, 0.009, and 0.0117 as well as RPCO-normalized data evaluated against CORUM protein complexes excluding cytoplasmic ribosomal gene pairs from the evaluation process.

